# ZYG-9^ch-TOG^ promotes the stability of acentrosomal poles via regulation of spindle microtubules in *C. elegans* oocyte meiosis

**DOI:** 10.1101/2022.01.04.474888

**Authors:** Timothy J. Mullen, Gabriel Cavin-Meza, Ian D. Wolff, Emily R. Czajkowski, Nikita S. Divekar, Justin D. Finkle, Sarah M. Wignall

**Affiliations:** Department of Molecular Biosciences, Northwestern University, Evanston, IL 60208

## Abstract

During mitosis, centrosomes serve as microtubule organizing centers that guide the formation of a bipolar spindle. However, oocytes of many species lack centrosomes; how meiotic spindles establish and maintain these acentrosomal poles remains poorly understood. Here, we show that the microtubule polymerase ZYG-9^ch-TOG^ is required to maintain acentrosomal pole integrity in *C. elegans* oocyte meiosis; following acute depletion of ZYG-9 from pre-formed spindles, the poles split apart and an unstable multipolar structure forms. Depletion of TAC-1, a protein known to interact with ZYG-9 in mitosis, caused loss of proper ZYG-9 localization and similar spindle phenotypes, further demonstrating that ZYG-9 is required for pole integrity. However, depletion of ZYG-9 surprisingly did not affect the assembly or stability of monopolar spindles, suggesting that ZYG-9 is not required for acentrosomal pole structure *per se*. Moreover, fluorescence recovery after photobleaching (FRAP) revealed that ZYG-9 turns over rapidly at acentrosomal poles, displaying similar turnover dynamics to tubulin itself, suggesting that ZYG-9 does not play a static structural role at poles. Together, these data support a global role for ZYG-9 in regulating the stability of bipolar spindles and demonstrate that the maintenance of acentrosomal poles requires factors beyond those acting to organize the pole structure itself.

## INTRODUCTION

When a cell divides, the genetic material must be accurately partitioned to ensure the viability of the daughter cells. This process is mediated by a bipolar microtubule-based spindle that provides the forces to segregate the chromosomes. In mitotically-dividing cells, two centriole-containing centrosomes nucleate microtubules and organize their minus ends to form the spindle poles, thus providing structural cues that impart stability to the spindle. Conversely, meiotically-dividing oocytes of many organisms lack centrosomes and therefore use distinct mechanisms to focus microtubule minus ends and arrange the spindle poles. Notably, acentrosomal spindle poles in human oocytes are frequently unstable; a significant fraction of oocytes undergo a period of spindle instability in which, following bipolar spindle formation, the poles split apart and come back together multiple times. This instability has adverse consequences, as oocytes that display more pole instability have higher rates of chromosome mis-segregation (Holubcova et al., 2015). However, despite the importance of organizing and stabilizing acentrosomal poles, little is known about what factors promote pole stability in oocytes of any organism.

Here, we use *C. elegans* as a model to address the question of how acentrosomal oocyte spindles are stabilized. The spindle assembly pathway has been well documented in this system. Microtubules are first nucleated and the minus ends are sorted outwards away from the chromosomes. The minus ends then organize into multiple nascent poles that coalesce until bipolarity is achieved (Wolff et al., 2016; Gigant et al., 2017). Importantly, this pathway is similar to what has been described in human oocytes, where multiple poles also form and then coalesce to form a bipolar spindle (Holubcova et al., 2015). A number of factors required for acentrosomal spindle assembly and pole formation in *C. elegans* have been identified: MEI-1/2^katanin^ promote the generation of short microtubules (Srayko et al., 2006), KLP-15/16^Kinesin-14^ bundle these microtubules (Mullen and Wignall, 2017), KLP-18^Kinesin-12^ and MESP-1 sort microtubules (Wolff et al., 2016; Wolff et al., 2021), ASPM-1 binds to microtubule minus ends and functions to focus the spindle poles in conjunction with dynein (Connolly et al., 2014; Cavin-Meza et al., 2021), and KLP-7^MCAK^ acts to limit the number of spindle poles and inhibit excess microtubule nucleation (Connolly et al., 2015; Han et al., 2015; Gigant et al., 2017). However, despite these proteins being well-studied in the context of spindle assembly, how the spindle is stabilized after it forms to maintain bipolarity is not well understood.

Now, we have gained insight into this question through studies of ZYG-9, a protein we have found to be required for the stability of acentrosomal spindles. In mitotically-dividing *C. elegans* embryos, ZYG-9 promotes the formation of long astral microtubules that aid in proper positioning of the mitotic spindle (Bellanger and Gonczy, 2003; Bellanger et al., 2007). In other organisms, ZYG-9 homologs (Stu2, Msps, XMAP215, ch-TOG) have also been shown to be essential for mitotic spindle assembly. In budding yeast, Stu2 is required for proper spindle positioning and orientation in addition to proper metaphase chromosome alignment (Kosco et al., 2001). In *Drosophila melangaster,* depletion of Msps in S2 cells results in shorter mitotic spindles (Goshima et al., 2005). In *Xenopus laevis* egg extracts, depletion of XMAP215 results in abnormally short spindles or failure of centrosomes to nucleate microtubules (Tournebize et al., 2000). Finally, in human cells, ch-TOG is necessary for spindle pole integrity, as ch-TOG depletion results in multipolar mitotic spindles (Gergely et al., 2003; Cassimeris and Morabito, 2004). Collectively, these phenotypes suggest that ZYG-9 family proteins regulate microtubule-based processes during cell division. This idea is also supported by *in vitro* experiments that demonstrate that ZYG-9-family proteins possess microtubule nucleation and polymerization activity (Brouhard et al., 2008; Widlund et al., 2011; Zanic et al., 2013; Thawani et al., 2018; Farmer et al., 2021).

In addition to this work on mitosis, prior studies have also implicated ZYG-9 family proteins as being essential for meiotic spindle assembly. Depletion of Msps results in multipolar spindles in *Drosophila* oocytes (Cullen and Ohkura, 2001). Moreover, in *C. elegans*, two early studies reported that spindles do not properly form following ZYG-9 depletion (Matthews et al., 1998; Yang et al., 2003), and a recent paper demonstrated that ZYG-9 is required for spindle pole coalescence (Chuang et al., 2020). However, whether ZYG-9 is required to maintain acentrosomal spindle stability once the bipolar spindle forms has not yet been investigated.

To address this question, we optimized and employed a degron-based strategy, enabling us to rapidly remove ZYG-9 from pre-formed bipolar spindles. This approach revealed that ZYG-9 is required to maintain the integrity of acentrosomal spindle poles; following ZYG-9 depletion the poles split apart and an unstable multipolar structure formed. Strikingly, this phenotype is similar to the unstable acentrosomal spindle poles described in human oocytes (Holubcova et al., 2015). However, despite its essential role in stabilizing bipolar spindles, we found that ZYG-9 depletion had no effect on monopolar spindle formation. This demonstrates that ZYG-9 is not required for pole integrity in all contexts; rather, the spindle destabilization seen following ZYG-9 depletion from pre-formed spindles likely arises from global effects on the spindle. Thus, our work has revealed new insight into mechanisms required for preserving meiotic spindle integrity and suggests that proper regulation of spindle microtubules is required to maintain the integrity of acentrosomal poles.

## RESULTS

### The XMAP215 homolog ZYG-9 is required for acentrosomal spindle pole coalescence

We previously performed an RNAi screen in *C. elegans* by knocking down proteins and screening oocytes arrested in Metaphase I for spindle defects (Wignall and Villeneuve, 2009; Mullen and Wignall, 2017). One hit from this screen was ZYG-9, a homolog of the microtubule polymerase XMAP215/ch-TOG. In *C. elegans* mitosis, depletion of ZYG-9 causes spindle positioning defects in the one-cell stage embryo due to shortened astral microtubule arrays, demonstrating a role for ZYG-9 in regulation of microtubule dynamics (Bellanger and Gonczy, 2003; Srayko et al., 2003). Moreover, depletion of ZYG-9 in oocytes has been reported to cause spindle defects (Matthews et al., 1998; Yang et al., 2003), but at the time we began our study, it was not known which stage of the acentrosomal spindle assembly pathway was affected. Therefore, to determine the cause of the metaphase defects, we depleted ZYG-9 using RNAi and filmed spindle assembly in oocytes expressing GFP::tubulin and mCherry::histone. In control oocytes, microtubules nucleate and then reorganize into an array with the minus ends oriented outwards; these ends subsequently form multiple poles that coalesce until bipolarity is achieved (Wolff et al., 2016). Following ZYG-9 depletion, microtubules nucleated and then the ends organized into multiple pole-like structures (Figure 1 – figure supplement 1A, Video 1); fixed imaging demonstrated that the microtubule minus end marker ASPM-1 and the spindle pole proteins MESP-1 and KLP-18 localized to these structures, confirming that they represent poles (Figure 1 – figure supplement 1B, 1C). However, the live imaging suggested that these poles were unstable, as we observed instances where multiple poles coalesced to form a transient bipolar-like spindle before splitting apart (Figure 1 – figure supplement 1A; Video 1). To quantify this phenotype we imaged single snapshots of intact worms, using oocyte position in the germ line as a proxy for meiotic progression. In *C. elegans*, oocytes are fertilized and then begin to form meiotic spindles as they pass through and exit the spermatheca into the “+1 position” of the germ line. Control embryos in the +1 position primarily contain either bipolar Meiosis I spindles (43%) or spindles that have progressed into Anaphase I (38%) and only 10% are still at the multipolar stage (Figure 1 – figure supplement 1D). In contrast, 50% of *zyg-9(RNAi)* embryos in the +1 position contained multipolar oocyte spindles while only 2% contained bipolar spindles, with the rest progressing to anaphase (Figure 1 – figure supplement 1D). These findings suggest that ZYG-9 is required for stable pole coalescence during acentrosomal spindle assembly, confirming the findings of a recent study (Chuang et al., 2020).

### A degron-based approach to investigate ZYG-9

The finding that acentrosomal poles transiently coalesce but then split apart following ZYG-9 depletion suggests that ZYG-9 may be required to maintain stable poles after they form. To test this hypothesis, we sought to rapidly deplete ZYG-9 from oocytes in which spindles had already established bipolarity. To this end, we took advantage of the auxin-inducible degron (AID) system (Zhang et al., 2015), using CRISPR to introduce a degron::GFP tag at the endogenous locus of *zyg-9* in a worm strain that expresses the TIR1 ubiquitin ligase from a germline-specific promoter (hereafter referred to as “ZYG-9 AID”; Figure 1A). This allows for proteasome-mediated degradation of degron-tagged ZYG-9 upon addition of the small molecule auxin.

**Figure 1:**
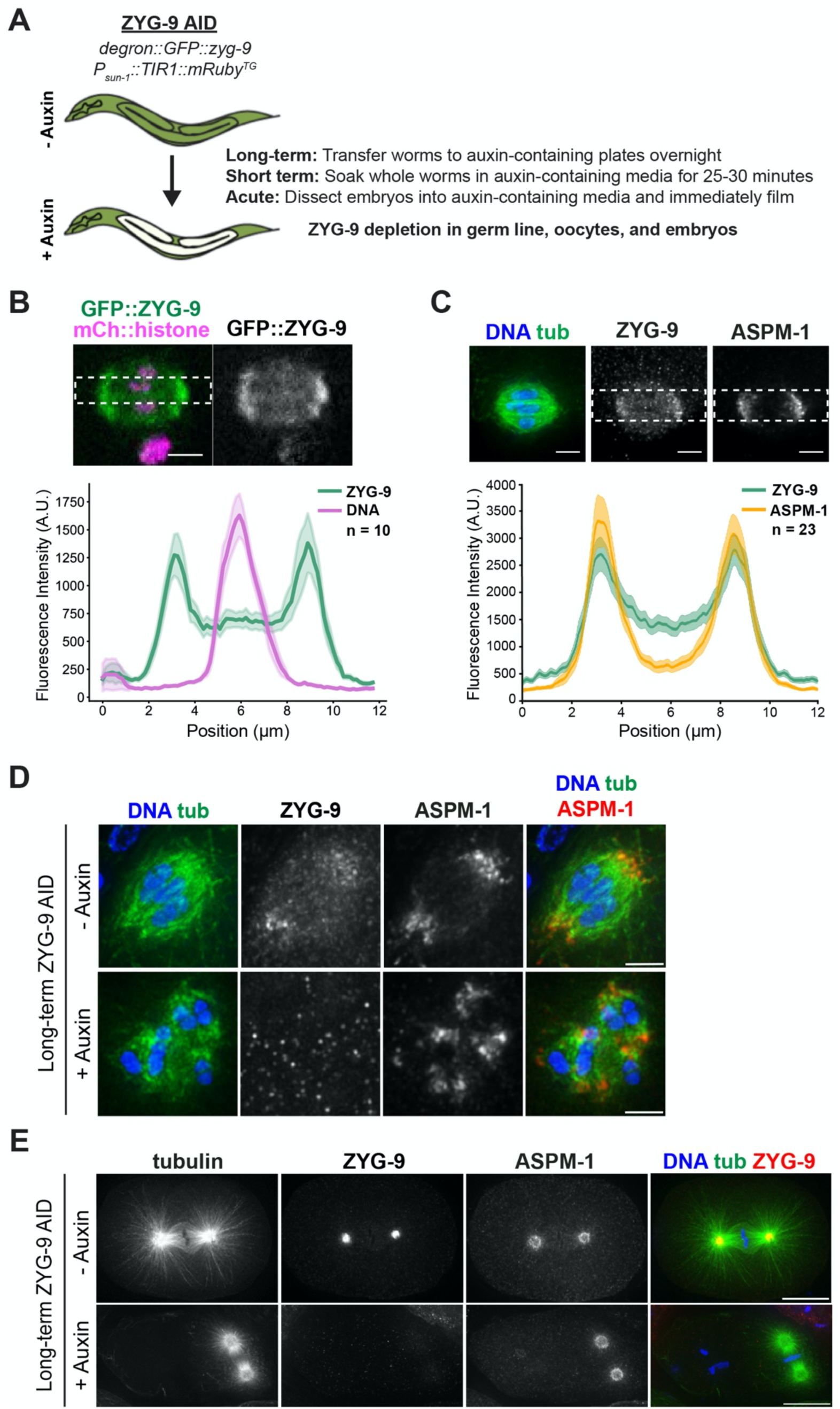
Validation of a ZYG-9-AID strain. (A) Schematic of the degron::GFP::ZYG-9 AID strain and description of auxin treatment lengths. ZYG-9 is tagged at the endogenous locus and the TIR1 transgene is expressed from a germline-specific promoter. Types of auxin treatment: (1) Long-term: worms are incubated on auxin-containing plates overnight, (2) Short-term: worms are soaked in auxin-containing media for 25-30 minutes prior to dissecting oocytes for immunofluorescence, (3) Acute: embryos are dissected into auxin-containing media and filmed immediately. (B) Linescan profiles of degron::GFP::ZYG-9 (green) and mCherry::histone (magenta) fluorescence intensities from metaphase spindles in live oocytes. The plot shows that ZYG-9 is enriched at the poles of the bipolar metaphase spindle, but ZYG-9 is localized in the midspindle region as well. n represents the number of spindles analyzed. Bar = 2.5 µm. (C) Linescan profiles of fixed metaphase spindles stained for DNA (blue), tubulin (green), degron::GFP::ZYG-9 (using the GFP antibody), and ASPM-1. The plot shows that ZYG-9 is enriched at the spindle poles but also has more signal in the midspindle region than ASPM-1. n represents the number of spindles analyzed. Bar = 2.5µm. (D) Meiotic oocyte spindles from ZYG-9 AID worms plated overnight on control plates (- auxin) and 1 mM auxin-containing plates (+ auxin) stained for DNA (blue), tubulin (green), ZYG-9 (not shown in merge), and ASPM-1 (red). Long-term ZYG-9 depletion results in the formation of multipolar spindles. Bars = 2.5µm. (E) Mitotic spindles in 1-cell stage embryos from ZYG-9 AID worms plated overnight on control plates (- auxin) and 1 mM auxin-containing plates (+ auxin) stained for DNA (blue), tubulin (green), ZYG-9 (red), and ASPM-1 (not shown in merge). Long-term ZYG-9 depletion recapitulates published mitotic phenotypes (Bellanger and Gonczy, 2003; Srayko et al., 2003). Bars = 10μm.

To confirm that degron::GFP:ZYG-9 retained its normal localization, we performed live imaging of oocytes and found that ZYG-9 begins to associate with microtubules at the multipolar stage, becoming enriched at the poles and increasing in intensity as the bipolar spindle forms (Figure 1 – figure supplement 2A, Figure 1B and Video 2), consistent with recent work (Chuang et al., 2020). Fixed imaging comparing the localization of degron::GFP::ZYG-9 to the spindle pole protein ASPM-1 confirmed these findings (Figure 1C, Figure 1 – figure supplement 2B). To further validate this strain, we incubated worms on auxin-containing plates overnight to mimic the long-term depletion achieved by RNAi. Under these conditions, the spindles appeared identical to those observed following *zyg-9(RNAi),* as they exhibited multiple ASPM-1-labeled poles (Figure 1D). Moreover, examination of mitotic spindles in 1-cell stage embryos after auxin treatment revealed misoriented mitotic spindles with shorter astral microtubules (Figure 1E), which phenocopies the mitotic defects reported for RNAi-mediated depletion of ZYG-9 (Bellanger and Gonczy, 2003; Srayko et al., 2003; Bellanger et al., 2007). Finally, we demonstrated that ZYG-9 was being depleted using both immunofluorescence (Figure 1D, 1E) and Western blotting (Figure 1 – figure supplement 3), confirming that our conditions could be used for efficient ZYG-9 depletion.

### ZYG-9 is required to maintain acentrosomal spindle pole integrity

After validating our ZYG-9 AID strain using long-term depletion, we next attempted to acutely deplete ZYG-9 and assess the stability of oocyte spindle poles that had already formed. To this end, we arrested oocytes at Metaphase I through RNAi-mediated depletion of EMB-30, a component of the anaphase promoting complex (Furuta et al., 2000), in a strain containing both degron::GFP::ZYG-9 and mCherry::tubulin. We then dissected these arrested oocytes into auxin-containing media and immediately mounted them for live imaging. Upon acute auxin treatment, we observed a rapid loss in GFP::ZYG-9 fluorescence, demonstrating that ZYG-9 depletion was occurring. Strikingly, as ZYG-9 was being depleted, the spindle poles began to split apart and fragment (Figure 2A and Videos 3 and 4). In contrast, oocytes dissected into media without auxin maintained bipolarity over the entire time course (Figure 2A, Video 5). These findings demonstrate that ZYG-9 is required to maintain acentrosomal pole stability.

**Figure 2:**
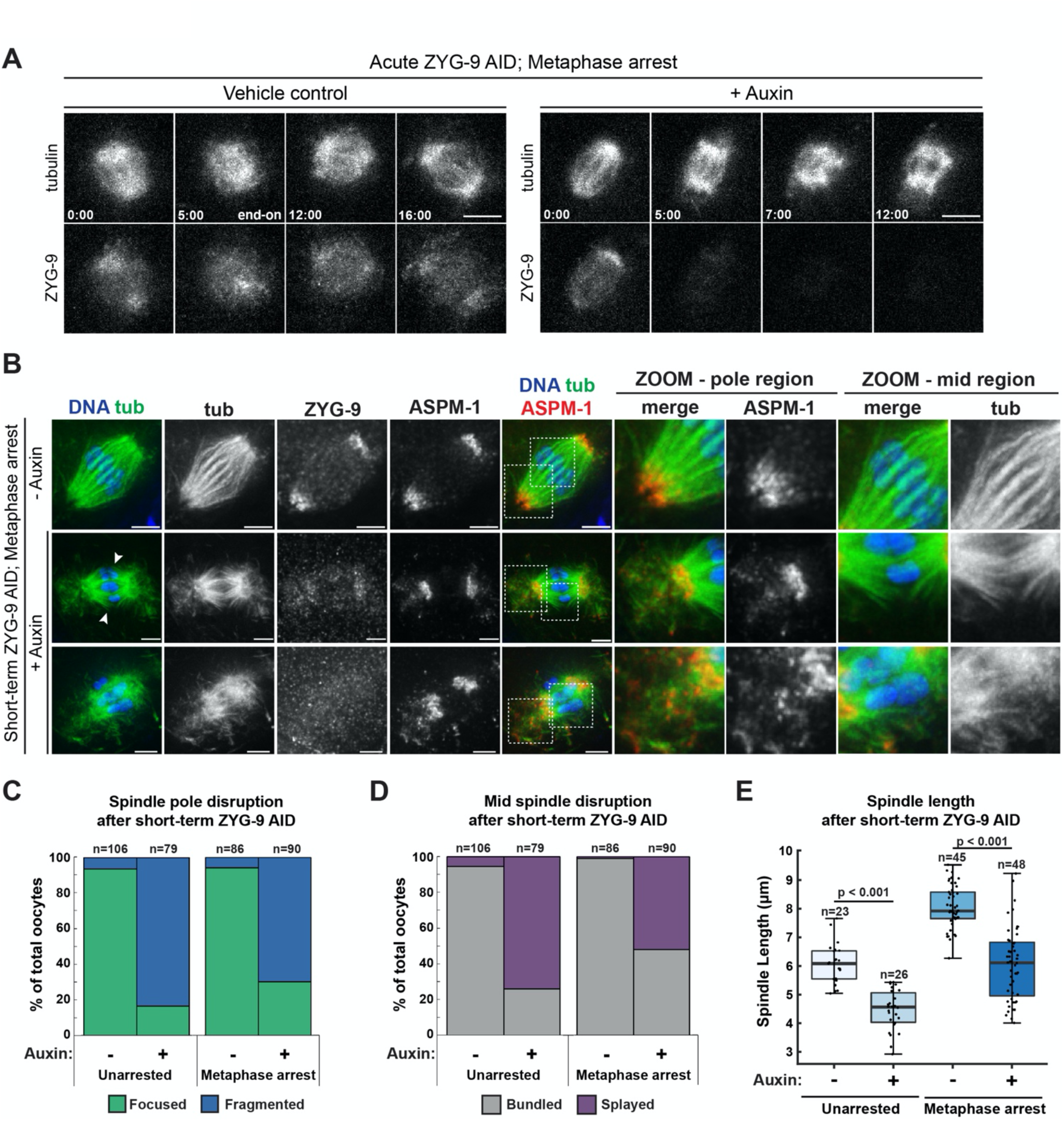
ZYG-9 is required to maintain spindle pole integrity. (A) Movie stills from Metaphase I-arrested *(emb-30(RNAi))* oocytes expressing mCherry::tubulin and degron::GFP::ZYG-9 acutely treated with either vehicle (left) or 100µM auxin (right). Following vehicle treatment, the oocyte spindle maintains ZYG-9 on the spindle and remains bipolar throughout the timecourse. Auxin treatment causes a rapid loss of ZYG-9 signal and a concurrent loss of spindle pole stability highlighted by the unfocusing and fragmentation of the spindle poles. Bars = 5 µm. Timestamp = min:sec. (B) Spindles from Metaphase I-arrested *(emb-30(RNAi))* oocytes treated with vehicle or 1mM auxin for 25-30 minutes. In vehicle-treated oocyte spindles, the poles are tightly organized, and the microtubule bundles appear to cross from one side of the spindle to the other without interruption. In spindles from auxin-treated oocytes, the ZYG-9 signal is decreased and spindle pole integrity is compromised as shown by the ASPM-1 and tubulin staining coming off the poles away from the spindle (zooms – pole region). Spindles from auxin-treated oocytes also showed defects in midspindle microtubules, where they appeared to terminate near the center of the spindle and splay away from the chromosomes (zooms – mid region). Bars = 2.5 µm. (C, D) Quantification of spindle pole phenotypes (C) or midspindle phenotypes (D) from unarrested (vector control) and Metaphase I-arrested *(emb-30(RNAi))* oocytes treated with either vehicle or auxin. n represents the number of spindles analyzed. (E) Quantification of spindle length (pole to pole) in unarrested and Metaphase I-arrested oocytes treated with either vehicle or auxin. Box represents the first quartile, median, and third quartile. Whiskers extend to maxima and minima. Significance determined using a two-tailed t-test. n represents the number of spindles analyzed.

To confirm this result and to examine ZYG-9-depleted spindles at higher resolution, we performed a short-term auxin treatment, soaking whole worms in auxin and subsequently dissecting Metaphase I-arrested oocytes and fixing them for immunofluorescence. Protein depletion using this fixed imaging method is less rapid than in dissected oocytes (Divekar et al., 2021b); we found that we needed to incubate worms for 25-30 minutes in auxin prior to dissection to begin to see spindle phenotypes. This is likely because auxin needs additional time to reach the oocytes when these cells are not dissected, resulting in increased incubation times and milder phenotypes. High resolution imaging of oocytes treated with auxin using this method revealed multiple pole defects, including unfocused spindle poles, microtubules emanating from the poles away from the chromosomes, and microtubules marked by ASPM-1 that appeared to be dissociated from the pole (Figure 2B, 2C), confirming that ZYG-9 is required to maintain the integrity of spindle poles. In addition to pole defects, we observed a large proportion of spindles that had defects in microtubule organization in the middle region of the spindle (Figure 2B arrowheads, 2D). In control oocytes, microtubule bundles run alongside chromosomes forming lateral associations (Wignall and Villeneuve, 2009). Although these bundles are thought to be comprised of many short microtubules based on electron microscopy (Srayko et al., 2006; Redemann et al., 2018), at the resolution of light microscopy the bundles appear to run continuously across the center of the spindle (Figure 2B, top row). In contrast, following ZYG-9 depletion, microtubule bundles appeared to splay away from the chromosomes (Figure 2B) and spindles were also significantly shorter than controls (Figure 2E). We observed similar phenotypes in oocytes that were not metaphase-arrested (Figure 2C, 2D, Figure 2 – figure supplement 1), demonstrating that these phenotypes were not caused by the metaphase arrest. Finally, we observed some spindles that exhibited defects, even though ZYG-9 was still detectable by immunofluorescence (Figure 2B, second row), suggesting that ZYG-9 does not have to be fully depleted to disrupt spindle organization. Together, these data suggest that ZYG-9 is continuously required to maintain the integrity of the oocyte spindle and is especially important for the stability of acentrosomal poles.

### The TACC homolog TAC-1 is required for proper ZYG-9 localization to the meiotic spindle

Next, we sought to investigate how ZYG-9 is regulated in oocyte meiosis. During mitosis, ZYG-9 forms a complex with the transforming acidic coiled-coil (TACC) homolog TAC-1 and these proteins are interdependent for localization to centrosomes (Bellanger and Gonczy, 2003; Srayko et al., 2003). To determine if TAC-1 also promotes the enrichment of ZYG-9 at acentrosomal poles, we imaged ZYG-9 localization in oocytes following *tac-1(RNAi)* (Figure 3A, 3B). ZYG-9 enrichment was lost at the poles in a majority of oocyte spindles observed (Figure 3C); we observed spindles where ZYG-9 was present on the spindle but no longer enriched at the poles (Figure 3A, row 2) as well as spindles where ZYG-9 localization was reduced overall (Figure 3A, rows 3-4). Moreover, there were spindle defects following *tac-1(RNAi)* that were similar to ZYG-9 depletion phenotypes, including fragmented poles with dispersed ASPM-1 (Figure 3A, 3B, 3D), disruption of midspindle microtubules (Figure 3A, 3B, 3E), and the formation of multipolar spindles (Figure 3A, row 4).

**Figure 3:**
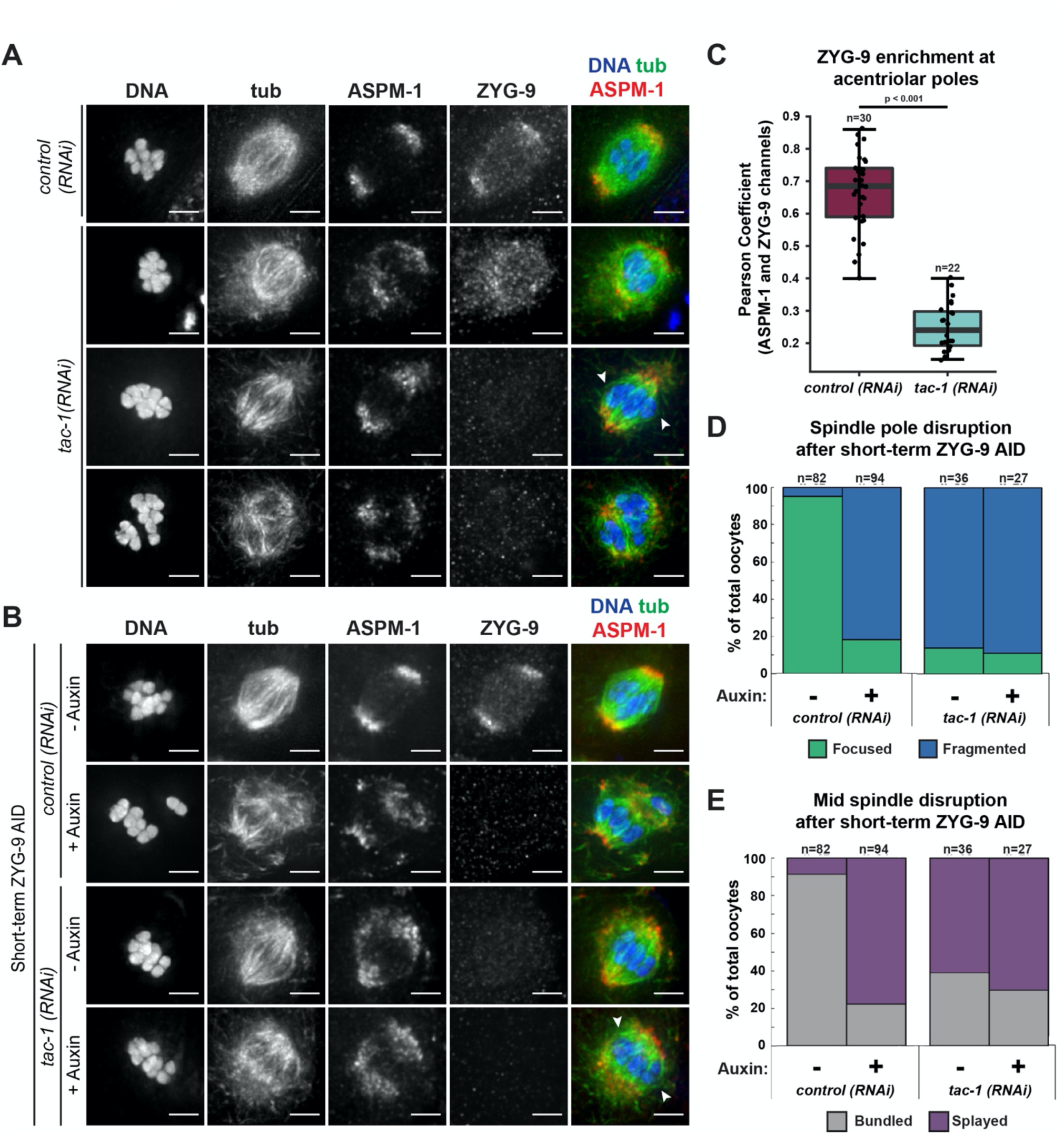
TAC-1 is required for proper localization of ZYG-9 to meiotic spindles. (A) IF imaging of oocyte spindles in either control or *tac-1(RNAi)* conditions. Shown are DNA (blue), tubulin (green), ASPM-1 (red), and ZYG-9 (not shown in merge). ZYG-9 enrichment at acentrosomal poles is lost in most spindles observed, and some spindles appear to have lost all ZYG-9 localization to the meiotic spindle; midspindle disruption is highlighted with arrowheads. Bars = 2.5µm. (B) IF imaging of oocyte spindles in either control or *tac-1(RNAi)* conditions. Shown are DNA (blue), tubulin (green), ASPM-1 (red), and ZYG-9 (not shown in merge). Whether subjected to *tac-1(RNAi)* alone or concurrently with short-term ZYG-9 AID depletion, spindle phenotypes mimic those observed in short-term ZYG-9 AID depletion alone (midspindle disruption is highlighted with arrowheads). Bars = 2.5µm. (C) Quantification of ZYG-9 enrichment at poles from oocyte spindles observed in (A); a Pearson coefficient was calculated for each image by comparing the ZYG-9 and ASPM-1 channels. Total spindles measured in each condition are noted above box plots; boxes represent first quartile, median, and third quartile. (D) Quantification of acentrosomal pole fragmentation from oocyte spindles observed in (B); total spindles counted in each condition are noted above stacked bars. (E) Quantification of midspindle microtubule splaying from oocyte spindles observed in (B); total spindles counted in each condition are noted above stacked bars.

Since we found that TAC-1 is required for robust ZYG-9 localization to the meiotic spindle and acentrosomal poles, we hypothesized that the *tac-1(RNAi)* phenotypes could be primarily explained by effects on ZYG-9. To test this, we performed short-term ZYG-9 AID depletion in worms treated with *tac-1(RNAi),* reasoning that if the spindle phenotypes worsened then it would suggest that TAC-1 has functions beyond recruiting ZYG-9 to the spindle. However, after treating *tac-1(RNAi)* worms with auxin, oocyte spindles appeared morphologically similar to either *tac-1(RNAi)* or ZYG-9 AID alone (Figure 3B). ASPM-1 labeling indicated clear pole fragmentation, and splaying of midspindle microtubules, and the percentages of pole fragmentation and midspindle disruption between these three depletion conditions were nearly identical (Figure 3D, 3E). Together, these data support the hypothesis that TAC-1 is functioning to properly localize ZYG-9 to the meiotic spindle and enrich ZYG-9 at acentrosomal poles.

Since TAC-1 and ZYG-9 are interdependent for localization to centrosomes in mitosis (Bellanger and Gonczy, 2003; Srayko et al., 2003), we next sought to perform the reciprocal experiment and assess TAC-1 localization in oocytes following ZYG-9 depletion. To this end, we first generated and validated a TAC-1 antibody (Figure 4 – figure supplement 1) and used it to assess TAC-1 localization throughout meiosis (Figure 4A). TAC-1 had no distinct localization on forming multipolar spindles, but once bipolarity was established, TAC-1 colocalized with ZYG-9 at acentrosomal poles and this colocalization persisted throughout anaphase. However, when we performed short-term ZYG-9 depletion from Metaphase I-arrested oocytes, TAC-1 localization to the spindle was lost (Figure 4B). These findings suggest that ZYG-9 and TAC-1 are interdependent for localization to both centrosome-containing and acentrosomal spindle poles.

**Figure 4:**
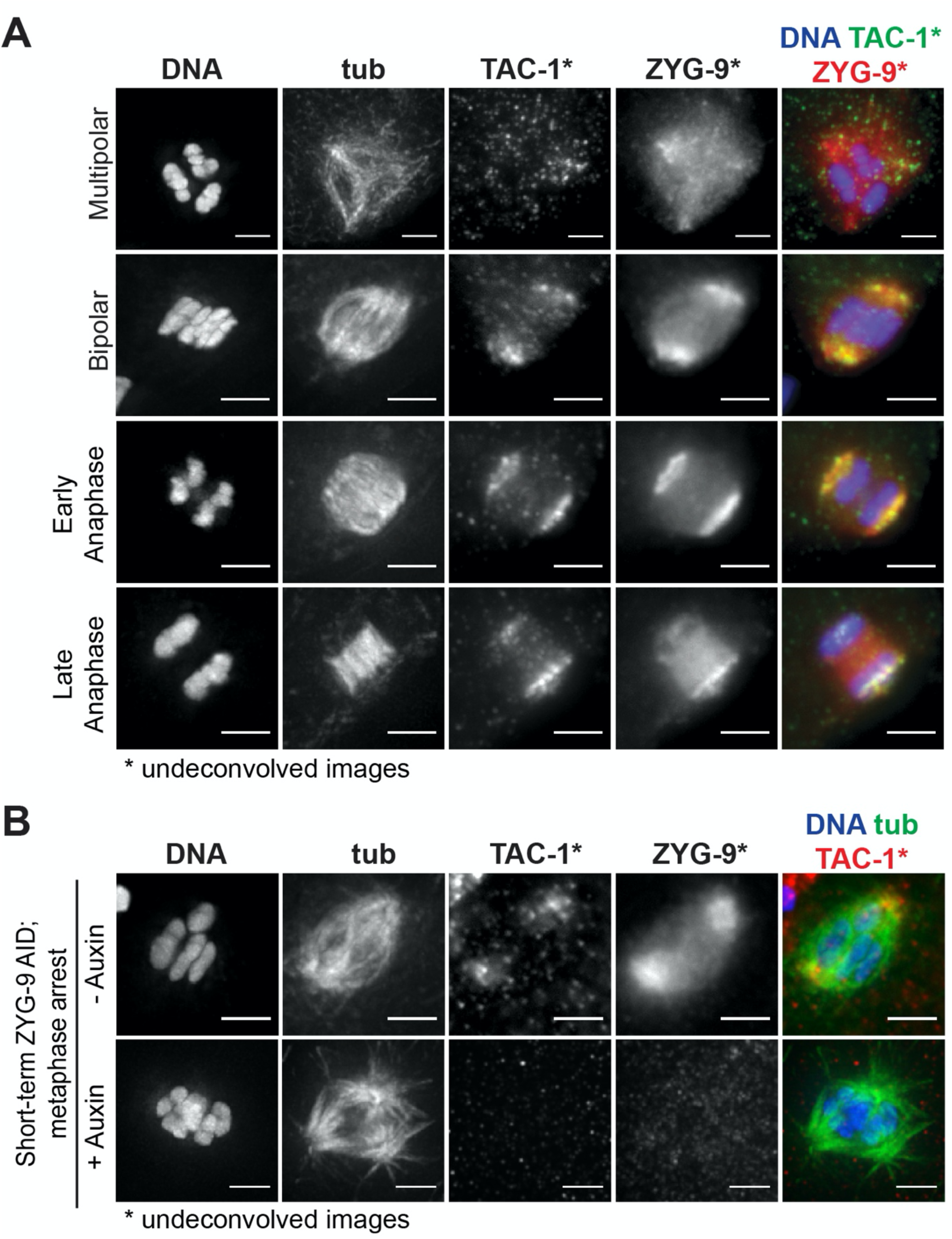
TAC-1 and ZYG-9 are interdependent for localization to acentrosomal poles. (A) IF imaging of oocyte spindles; shown are DNA (blue), TAC-1 (green), ZYG-9 (red), and tubulin (not shown in merge). Colocalization of TAC-1 and ZYG-9 is evident in metaphase and persists throughout anaphase. DNA and Tubulin channels were deconvolved, while ZYG-9 and TAC-1 channels were not due to higher background staining of the TAC-1 antibody. Bars = 2.5µm. (B) IF imaging of oocyte spindles in Metaphase I-arrest *(emb-30(RNAi))* conditions; shown are DNA (blue), tubulin (green), TAC-1 (red), and ZYG-9 (not shown in merge). Short-term ZYG-9 depletion disrupts localization of TAC-1 to acentrosomal poles. As in (A), ZYG-9 and TAC-1 channels were not deconvolved. Bars = 2.5µm.

### ZYG-9 is not required to form or stabilize spindle poles in all contexts

Next, we sought to investigate how ZYG-9 promotes acentrosomal spindle stability. Since ZYG-9 localizes to spindle poles (Figure 1B, 1C, Figure 1 – figure supplement 2) and is required for both pole coalescence (Chuang et al., 2020) (Figure 1 – figure supplement 1, Figure 1D) and maintenance (Figure 2A-C, Figure 2 – figure supplement 1), one possibility is that a population of ZYG-9 at spindle poles functions to organize microtubule minus ends into a stable structure. However, since ZYG-9 depletion also causes microtubule bundles in the center of the spindle to splay (Figure 2B, 2D, Figure 2 – figure supplement 1) and decreases spindle length (Figure 2E), it is possible that the pole defects could instead be the result of a more global effect on spindle microtubules. For instance, if ZYG-9 depletion affects microtubules in the center of the spindle, disruption of this region could destabilize the entire structure and lead to pole splitting. Consistent with this second possibility, we observed a substantial population of ZYG-9 in the center of the spindle (Figure 1B). Notably, not all spindle pole components have this midspindle population, as the intensity of ASPM-1 fluorescence is much less prominent in this region (Figure 1C).

Consequently, to explore these possibilities we asked whether ZYG-9 depletion would affect the formation of monopolar spindles. These structures have a single pole and therefore lack the overlap region that is present at the center of bipolar spindles. Thus, if monopoles are disrupted following ZYG-9 depletion, it would suggest that ZYG-9 has a structural role at poles, rather than the pole defects in bipolar spindles arising through disruption of the central overlap zone. To generate monopolar spindles, we depleted KLP-18^kinesin-12^ via RNAi in the ZYG-9 AID strain; in oocytes lacking KLP-18, microtubule minus ends fail to be sorted outwards during spindle assembly and instead organize into a single ASPM-1-marked pole (Wignall and Villeneuve, 2009; Wolff et al., 2016). We then performed short-term ZYG-9 depletion by soaking *klp-18(RNAi)* worms in auxin, followed by immunofluorescence to assess oocyte spindle morphology. Importantly, monopolar spindles appeared normal following ZYG-9 depletion; there was a single, ASPM-1-marked pole and the volume of ASPM-1 at the pole was indistinguishable in the presence or absence of ZYG-9 (Figure 5A, 5B). As a complementary approach, we performed live imaging to visualize the effects of acute ZYG-9 removal from pre-formed monopolar spindles. Upon auxin treatment, we observed a rapid loss in degron::GFP::ZYG-9 fluorescence but this depletion did not cause any noticeable effects on the stability or organization of the monopole; ZYG-9-depleted monopolar spindles maintained their aster-like structure, similar to control spindles where ZYG-9 was present (Figure 5C, Videos 6, 7). Thus, ZYG-9 is not required to maintain acentrosomal pole structure, and instead performs a function that is specifically required for the formation and stabilization of bipolar spindles. This result, in combination with the midspindle defects observed following ZYG-9 depletion (Figure 2B, 2D, Figure 2 – figure supplement 1), suggests that ZYG-9 does not solely function at acentrosomal spindle poles, and instead may promote pole stability through more global effects on spindle microtubules.

**Figure 5:**
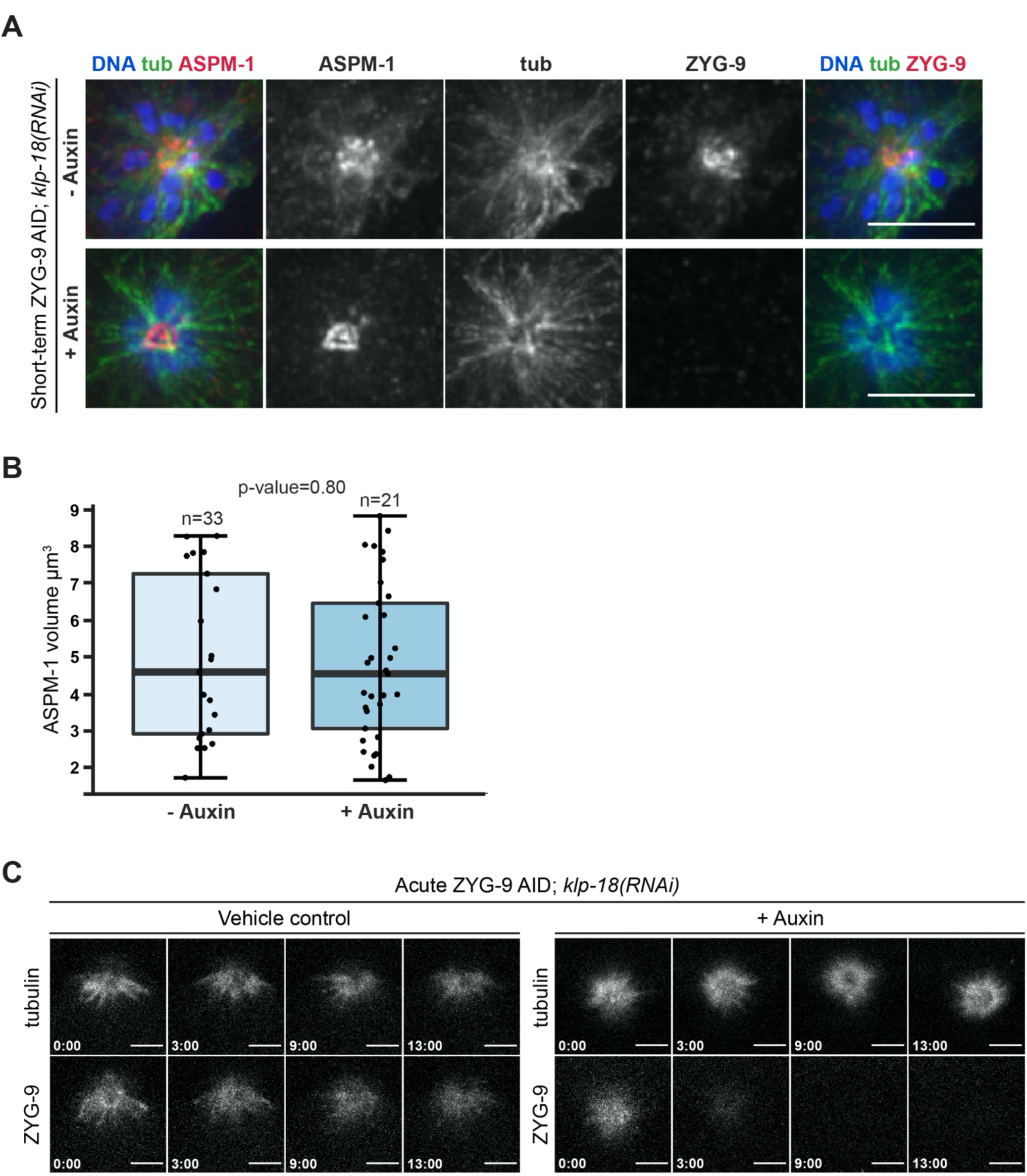
ZYG-9 is not required for acentrosomal pole stability of monopolar spindles. (A) (B) IF imaging of oocyte spindles in *klp-18(RNAi)* conditions; shown are DNA (blue), microtubules (green), ZYG-9 (red, merge on the right), and ASPM-1 (red, merge on the left). When depleting ZYG-9 via short-term auxin treatment, monopolar spindles appear unperturbed, as can be seen through a singular, focused pole of ASPM-1. Bars = 2.5µm. (B) Quantification of IF imaging represented in (A); there is no significant difference in the volume of monopolar spindles between untreated and auxin-treated conditions. Box represents the first quartile, median, and third quartile. Whiskers extend to maxima and minima. Significance determined using a two-tailed t-test. n represents the number of spindles analyzed. (C) Movie stills from *klp-18(RNAi))* oocytes expressing mCherry::tubulin and degron::GFP::ZYG-9 acutely treated with either vehicle (left) or 100µM auxin (right). Auxin treatment causes a rapid loss of ZYG-9 signal but the structure of the monopolar spindle does not noticeably change. Bars = 5 µm. Timestamp = min:sec.

### ZYG-9 is highly dynamic at acentrosomal spindle poles

While centrosomes act as structural cues that organize mitotic spindle poles, it is not clear whether there are factors that stably associate with acentrosomal poles and perform this scaffolding role in *C. elegans* oocytes. Given that our data suggest that ZYG-9 stabilizes poles through a global effect on spindle microtubules, we would not expect it to act as this type of static scaffold. However, since nothing is known about the dynamics of acentrosomal pole proteins in *C. elegans*, we set out to test this prediction directly.

A previous study assessed ZYG-9 dynamics at the poles of mitotic centrosome-containing spindles using fluorescence recovery after photobleaching (FRAP) (Woodruff et al., 2017). That investigation demonstrated that the exchange of ZYG-9 between the centrosome and cytoplasm was not highly dynamic, suggesting that it is a relatively stable component of centrosome-containing poles. To determine if this is also true at acentrosomal spindle poles, we performed an analogous experiment by photobleaching GFP::ZYG-9 and assessing its recovery at the poles of oocyte spindles compared to at centrosomes (Figure 6A, Figure 6 – figure supplement 1A). To assess the stability of ZYG-9 in mitosis we imaged centrosomes in the EMS (endomesodermal precursor) cell of 4-cell embryos since these cells remain stable at metaphase for a few minutes, allowing us to image fluorescence recovery (Decker et al., 2011; Woodruff et al., 2017). This experiment revealed that ZYG-9 turns over much more rapidly at acentrosomal spindle poles (t_1/2_ = 16.61s) compared to centrosomes (t_1/2_ = 119.39s) during metaphase (Figure 6A and Figure 6 – figure supplement 1A; Videos 8 and 9). Furthermore, ZYG-9 fluorescence at acentrosomal poles recovered to pre-bleach levels during the time course whereas ZYG-9 fluorescence at centrosomes did not. Strikingly, the rate of ZYG-9 turnover at acentrosomal poles was so rapid that it was similar to the recovery of tubulin (t_1/2_ = 14.61s) (Figure 6B and Figure 6 – figure supplement 1B; Video 10). These results suggest that ZYG-9 is not a stable component of acentrosomal poles.

**Figure 6:**
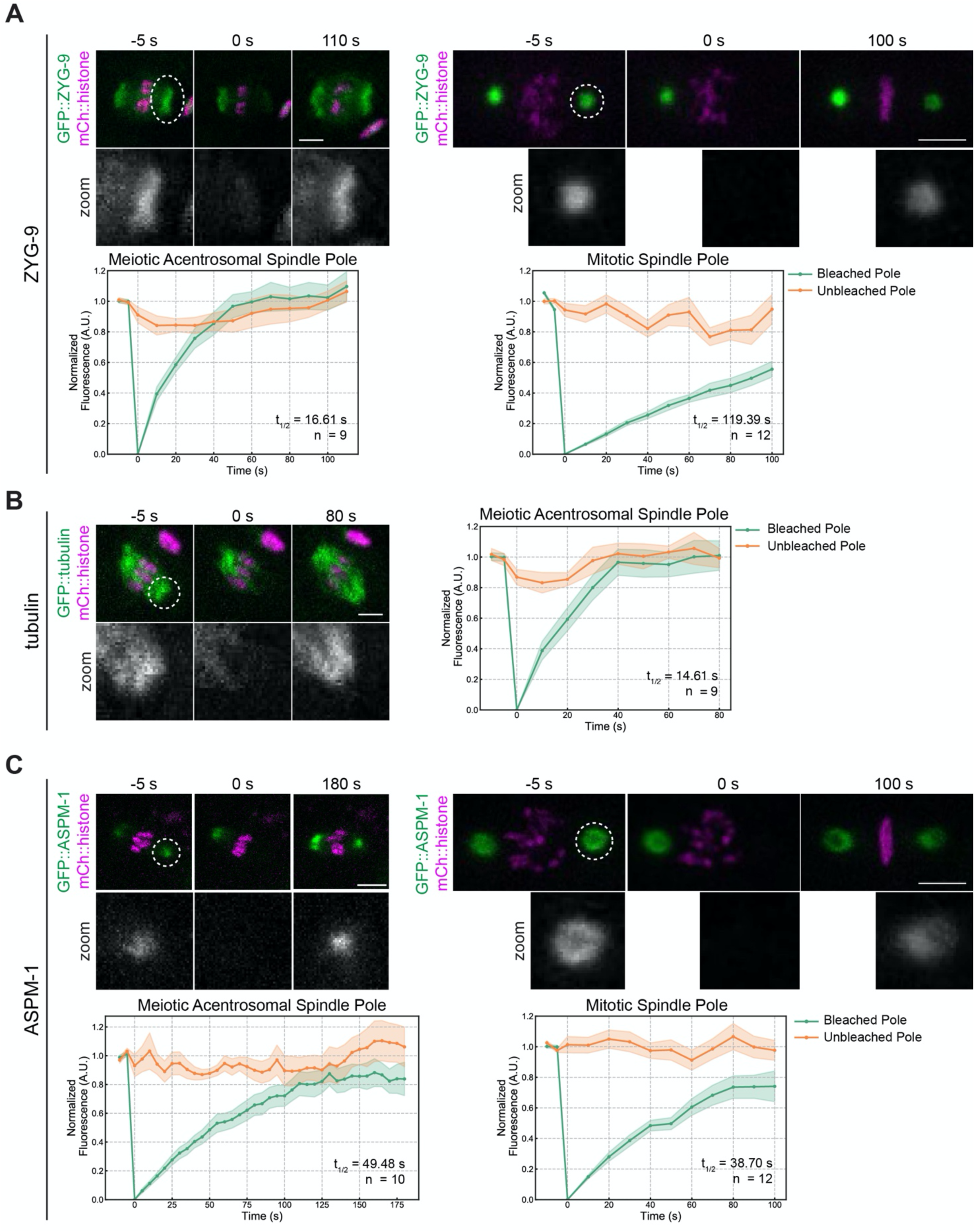
FRAP analysis of ZYG-9 and ASPM-1 at acentrosomal poles compared to at centrosomes. (A-C) FRAP recovery curves and stills from movies for ZYG-9 (A), tubulin (B), and ASPM-1 (C) at meiotic and mitotic spindle poles (except for tubulin; only recovery at the meiotic spindle is shown). In graphs, the bleached pole curve is green, the unbleached pole curve is orange, the solid lines are the average, and the standard error of the mean is shaded. For all images, the specified bleached protein is in green and the chromosomes are in magenta. Zooms show only the bleached region. t1/2 was calculated from fitting the recovery curves to a single exponential function (Materials and Methods); see Supplemental Figure 5. n represents the number of spindles analyzed to generate each curve. ZYG-9 is highly dynamic at acentrosomal spindle poles, displaying a similar recovery time to tubulin, while ASPM-1 turns over less rapidly and has similar dynamics at centrosome-containing and acentrosomal poles. Bars = 2.5μm (meiosis) and 5μm (mitosis).

Following on these findings, we sought to investigate whether the faster ZYG-9 turnover in oocytes was the result of the inherent organization of acentrosomal poles, rather than reflecting a specific difference in ZYG-9 behavior; if this were the case, other pole components would also show the same discrepancy in dynamics compared to centrosomes. To test this, we performed FRAP analysis on ASPM-1, and found that this protein had relatively similar turnover rates both in oocytes and in mitotically-dividing embryos (t_1/2_ = 49.48s and 38.70s, respectively) and the fluorescence recovered to similar levels during the time of filming (Figure 6C and Figure 6 – figure supplement 1C; Videos 11 and 12). Moreover, ASPM-1 recovered more slowly than ZYG-9 and tubulin (Figure 6A-C, Figure 6 – figure supplement 1), demonstrating that not all pole proteins display the rapid turnover seen for ZYG-9 at acentrosomal poles. Therefore, the rapid turnover of ZYG-9 in oocytes is not solely due to differences in pole organization in the absence of centrosomes. Rather, the faster ZYG-9 dynamics imply that ZYG-9 does not play a static, structural role at acentrosomal poles, and instead support the idea that ZYG-9 acts more globally to regulate spindle stability during oocyte meiosis.

## DISCUSSION

### ZYG-9 is required to stabilize and maintain acentrosomal poles during oocyte meiosis

Taken together, our data support a model in which ZYG-9 provides spindle stability throughout meiosis (Figure 7). ZYG-9 begins to localize to the spindle as multiple poles are formed; these poles then merge, forming a stable bipolar structure. Without ZYG-9 present, nascent poles marked by ASPM-1 form, but can never stably coalesce. This finding was also recently published by another group, who depleted ZYG-9 in *C. elegans* oocytes using RNAi and reported similar depletion phenotypes as those seen with long-term ZYG-9 depletion in our degron system (Chuang et al., 2020). Our current study now extends these findings, demonstrating that ZYG-9 is also required to maintain spindle stability. We found that when ZYG-9 was removed from a pre-formed spindle, acentrosomal poles destabilize and the overlap zone of the spindle is disrupted, causing the spindle to become unstable and ultimately lose bipolarity. Thus, ZYG-9 is not only required for spindle assembly but is also continuously required to maintain spindle bipolarity.

**Figure 7:**
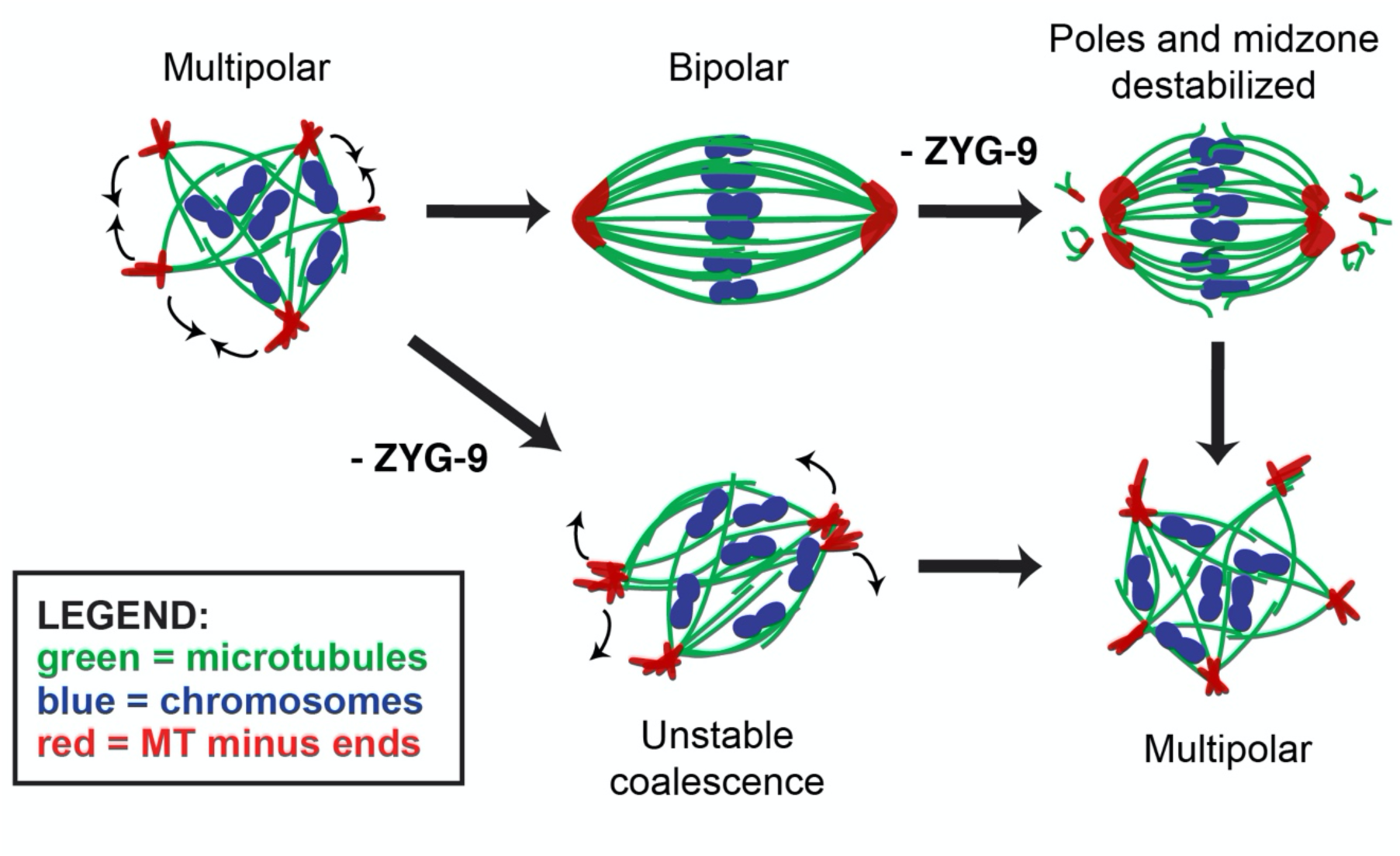
Model of acentrosomal pole coalescence and stability provided by ZYG-9. ZYG-9 is required to establish and maintain acentrosomal pole stability during meiosis. Removal of ZYG-9 prior to spindle assembly prevents multipolar spindles from stably coalescing into bipolar spindles. Additionally, removing ZYG-9 from stable bipolar spindles causes severe splaying of microtubule bundles near chromosomes and fragmentation of acentrosomal poles, ultimately causing the spindle to lose bipolarity and revert to a multipolar state, demonstrating that ZYG-9 is continuously required for maintaining stable bipolar spindles.

Intriguingly, the observation that ZYG-9 depletion does not affect monopolar spindle pole structure demonstrates that the pole fragmentation phenotype does not occur in all contexts. This suggests that ZYG-9 is not required for pole formation *per se*, and instead regulates spindle stability more globally. We speculate that ZYG-9 could perform this function by regulating microtubule dynamics. The *Xenopus* homolog of ZYG-9, XMAP215, has been extensively characterized, and is known to be able to promote microtubule nucleation through an interaction with the ɣ-tubulin ring complex (Thawani et al., 2018). Moreover, XMAP215 has been demonstrated to have microtubule polymerase activity; this protein utilizes multiple TOG domains to bind tubulin dimers for incorporation into growing microtubule plus ends (Gard and Kirschner, 1987; Brouhard et al., 2008; Widlund et al., 2011; Ayaz et al., 2012). Additionally, recent work has demonstrated that XMAP215 can also promote microtubule catastrophe (Farmer et al., 2021), implicating this protein in regulating multiple aspects of microtubule dynamics. In *C. elegans*, *in vitro* work with ZYG-9 has revealed that a minimal condensate comprised of ZYG-9, the microtubule associated protein TPXL-1, and the centrosomal protein SPD-5 can generate microtubule asters with sufficient tubulin concentrations (Woodruff et al., 2017). Finally, a role in regulating microtubule dynamics is consistent with the *in vivo* phenotypes of ZYG-9 depletion; astral microtubules are substantially shorter following ZYG-9 depletion in mitotically-dividing embryos (Matthews et al., 1998; Bellanger and Gonczy, 2003; Srayko et al., 2003), and we demonstrated in this study that oocyte spindles shorten following ZYG-9 depletion, before they lose bipolarity. Altogether, this evidence makes it likely that ZYG-9 regulates microtubule dynamics *in vivo*. One possibility is that ZYG-9, by acting as a microtubule polymerase, creates the appropriate microtubule substrate for sorting factors like KLP-18 and MESP-1. These factors sort bundles outwards where their ends can then be coalesced into stable poles; without ZYG-9 these proteins would therefore not be able to perform their proper functions, resulting in disruption of spindle organization. However, it is also possible that ZYG-9 facilitates microtubule nucleation and/or promotes catastrophe, as has been shown for XMAP215. Further investigation will be invaluable for evaluating how ZYG-9 affects microtubule dynamics in *C. elegans* oocytes, to better understand the role of ZYG-9 in the maintenance of spindle stability.

### TAC-1 is essential for proper ZYG-9 localization to acentrosomal spindles

In addition to gaining insight into the role of ZYG-9, we also demonstrated that ZYG-9 and TAC-1 are interdependent for proper localization to the meiotic spindle, similar to what has been observed in *C. elegans* mitotically-dividing embryos (Bellanger and Gonczy, 2003; Srayko et al., 2003). Depletion of TAC-1 via RNAi leads to multiple phenotypes with varying degrees of acentrosomal pole instability, and concurrent depletion of ZYG-9 and TAC-1 does not exacerbate the spindle phenotypes when compared to ZYG-9 AID depletions alone. These data suggest that TAC-1 functions to localize ZYG-9 to meiotic spindle poles, but it remains unclear how this interaction is regulated. In *Xenopus* egg extracts, the TACC homolog Maskin is phosphorylated by Eg2 (Aurora A kinase) and this phosphorylation regulates Maskin’s localization and function (Peset et al., 2005). Related studies of mitotic *Drosophila* embryos have demonstrated that Aurora A stabilizes the interaction between TACC and XMAP215 homologs (D-TACC and Msps, respectively) (Barros et al., 2005). The *C. elegans* Aurora A ortholog, AIR-1, is essential for spindle assembly and microtubule nucleation in oocytes (Sumiyoshi et al., 2015), raising the possibility that it could work in concert with ZYG-9. However, whether Aurora A regulates the interaction between TAC-1 and ZYG-9 in *C. elegans* oocytes remains to be investigated.

During mitosis, ZYG-9 and TAC-1 have also been shown to form a complex with another kinase, ZYG-8 (Bellanger et al., 2007). ZYG-8 mutants were shown to have similar mitotic defects as those seen with ZYG-9 and TAC-1 RNAi (Gonczy et al., 2001; Bellanger et al., 2007), but the function of ZYG-8 during oocyte meiosis has not been extensively characterized. A previous study observed defects in anaphase spindle elongation with removal of ZYG-8 activity via a temperature-sensitive mutation, suggesting a decrease in microtubule polymerization (McNally et al., 2016), but no spindle assembly or stability phenotypes were noted prior to anaphase. It is possible that ZYG-8 is responsible for phosphorylating TAC-1 in *C. elegans* oocyte meiosis to stabilize the interaction between TAC-1 and ZYG-9, rather than Aurora A. If phosphorylation of TAC-1 by ZYG-8 stabilizes the interaction of TAC-1 and ZYG-9 and their localization to the meiotic spindle, loss of ZYG-8 activity could lead to loss of ZYG-9 polymerase activity, which could account for the reduced anaphase spindle elongation previously observed. More detailed observations of ZYG-8 localization and loss of ZYG-8 activity prior to anaphase will help to determine if ZYG-8 is interacting with ZYG-9/TAC-1 and will clarify its role in oocyte meiosis.

### Acentrosomal pole proteins have distinct and separate contributions to pole coalescence and stability

To date, the mechanisms of microtubule nucleation and polymerization during meiotic acentrosomal spindle assembly in *C. elegans* are not well characterized. Recent work in this system has implicated the nuclear lamina, the Ran GTPase gradient, and ɣ-tubulin in microtubule nucleation (Chuang et al., 2020). Additionally, the work presented in both Chuang *et al*. and this study supports a role for ZYG-9 in promoting microtubule polymerization and suggests that proper regulation of microtubule dynamics may be essential for spindle pole coalescence and for the maintenance of acentrosomal spindle bipolarity. Building on these findings, it will be important to define the exact hierarchal structure of proteins that act to assemble the pole and then to coalesce and stabilize these structures. While localization of numerous proteins to acentrosomal poles has been reported (reviewed in (Severson et al., 2016; Mullen et al., 2019)), how each protein contributes to pole coalescence and maintenance remains a crucial gap in knowledge.

Through FRAP of acentrosomal spindle pole components, we have begun to tease apart differences in the dynamics of pole proteins, providing some insight into their functions. Notably, ZYG-9 recovered from bleaching in the same timeframe as tubulin itself, suggesting that ZYG-9 is extremely dynamic on the acentrosomal spindle. This is in contrast to another pole protein, ASPM-1, which took longer to recover. ASPM-1 localization to meiotic spindle poles is robust and difficult to disrupt; this protein is often used as a marker for poles following depletions of various other spindle proteins. Additionally, ASPM-1 has been shown to be required for localizing other pole proteins, such as LIN-5 and dynein (van der Voet et al., 2009). The slower turnover for ASPM-1 raises the possibility that it could be crucial for building an initial pole structure, on which other pole proteins can act upon to drive coalescence and stability. Notably, pole splaying has been observed following RNAi of ASPM-1 or dynein/dynactin (Yang et al., 2005; Ellefson and McNally, 2009; van der Voet et al., 2009; Ellefson and McNally, 2011; Crowder et al., 2015; Muscat et al., 2015), supporting a role for these proteins in pole organization.

In a recent study, we used the degron system to investigate dynein, and found that, like ZYG-9, dynein is required for both pole organization and stability (Cavin-Meza et al., 2021). However, the phenotype of dynein depletion was markedly different from what we report for ZYG-9 depletion in the current study; while acute dynein depletion caused focused poles to splay, the center of the spindle remained intact and therefore the spindles were able to maintain bipolarity. Moreover, when we depleted dynein from monopolar spindles, the entire monopole came apart, dispersing microtubule bundles into the cytoplasm. These findings suggest that in contrast to ZYG-9, dynein plays a specific role at the pole itself, acting to organize microtubule minus ends into a stable pole structure. In contrast, our findings with ZYG-9 and TAC-1 suggest that ZYG-9 has a distinct, broader function in meiotic spindles that is not restricted to keeping acentrosomal poles focused and stable.

In summary, our degron-based experiments have provided new insight into the role of ZYG-9 in stabilizing the acentrosomal spindle and suggest that proper regulation of microtubule dynamics is essential to maintain spindle pole integrity. Applying degron-based approaches to other pole proteins and comparing the depletion phenotypes on bipolar and monopolar spindles, will help elucidate the contributions of other pole components to spindle assembly and/or maintenance. Intriguingly, the spindle instability observed in our live imaging of acute ZYG-9 depletion is reminiscent of the instability that has been observed in human oocytes (Holubcova et al., 2015). Loss of bipolarity, as seen following ZYG-9 depletion, would lead to abnormal segregation of chromosomes into multiple masses; this is similar to the correlation between spindle instability and chromosome segregation errors seen in human oocytes (Holubcova et al., 2015). Therefore, it would be exciting for future studies to investigate whether misregulation of microtubule dynamics could contribute to the spindle instability seen in human oocytes.

## MATERIALS AND METHODS

### Strains

EU1067: *unc-119(ed3) ruIs32 [unc-119(+) pie-1::GFP::H2B]* III; *ruIs57 [unc-119(+) pie-1::GFP::tubulin] (*gift from Bruce Bowerman)

OD56: *ltIs37 [(pAA64) pie-1::mCherry::his-58 + unc-119(+)] IV* (gift from Arshad Desai)

OD57: *unc-119(ed3)* III; *ltIs37 [pAA64; pie-1::mCherry::his-58; unc-119(+)]* IV; *ltIs25 [pAZ132; pie-1::GFP::tba-2; unc-119 (+)]* (gift from Arshad Desai)

EU2876: *or1935 [GFP::aspm-1] I; ltIs37 [pAA64; pie-1::mCherry::his-58; unc-119(+)] IV* CA1199: *unc-119(ed3); ieSi38 [Psun-1::TIR1::mRuby::sun-1 3’UTR, cb-unc-119(+)] IV* (gift from Abby Dernburg)

JA1559: *weIs21 [pJA138 (pie-1::mCherry::tub::pie-1)]; unc-119(ed3) III* (gift from Mike Glotzer) SMW21: CA1199 x OD56; *unc-119(ed3); ieSi38 [Psun-1::TIR1::mRuby::sun-1 3’UTR, cb-unc-119(+)] IV; ltIs37 [(pAA64) pie-1::mCherry::his-58 + unc-119(+)] IV*

SMW22: CA1199 x JA1559; *unc-119(ed3); ieSi38 [Psun-1::TIR1::mRuby::sun-1 3’UTR, cb-unc-119(+)] IV; weIs21 [pJA138 (pie-1::mCherry::tub::pie-1)]; unc-119(ed3) III*

SMW24: *Pzyg-9::degron::EmGFP::zyg-9 (C6->T—PAM site mutation) II; unc-119(ed3); ieSi38 [Psun-1::TIR1::mRuby::sun-1 3’UTR, cb-unc-119(+)] IV*

SMW26: SMW24 x SMW21; *Pzyg-9::degron::EmGFP::zyg-9 (C6->T—PAM site mutation) II; unc-119(ed3); ieSi38 [Psun-1::TIR1::mRuby::sun-1 3’UTR, cb-unc-119(+)] IV; ltIs37 [(pAA64) pie-1::mCherry::his-58 + unc-119(+)] IV*

SMW33: SMW24 x SMW22; *Pzyg-9::degron::EmGFP::zyg-9 (C6->T—PAM site mutation) II; unc-119(ed3); ieSi38 [Psun-1::TIR1::mRuby::sun-1 3’UTR, cb-unc-119(+)] IV; weIs21 [pJA138 (pie-1::mCherry::tub::pie-1)]; unc-119(ed3) III*

### Generation of degron::EmGFP::ZYG-9 strain (SMW24)

A CRISPR-based approach (Arribere et al., 2014; Paix et al., 2015) was used to generate an endogenously tagged degron::EmGFP::ZYG-9 (SMW24). Briefly, 27μM recombinant Alt-R *S. pyogenes* Cas9 protein (IDT) was co-injected with 13.6μM tracrRNA (IDT), 4μM *dpy-10* crRNA, 1.34μM *dpy-10* repair oligo, 9.6μM *zyg-9* crRNA, and 136ng/μL ssDNA *zyg-9* repair template into CA1199 (Psun-1::TIR1::mRuby) worms, that were then allowed to produce progeny (See Table 1 for list of tracrRNA, crRNA, and primers used). Worms from plates containing rollers and dumpys were screened for degron::GFP insertions by PCR screening. To make the *zyg-9* repair template, we generated an N-terminal degron::EmGFP::linker (pADR28) using site directed mutagenesis and pLZ29 (gift from Abby Dernburg). The linker we inserted is from pIC26. The tag was then amplified using PCR with primers that contained homology to the *zyg-9* locus with the final product containing 57 bp of homology upstream of the *zyg-9* start codon and 61 bp of homology downstream of the start codon. ssDNA was generated by asymmetric PCR. SMW24 (degron::EmGFP::ZYG-9) was crossed with SMW21 (mCherry::histone; TIR1::mRuby) to generate SMW26: *Pzyg-9-16::degron::EmGFP::ZYG-9 (C6->T—PAM site mutation) II; ieSi38 [Psun-1::TIR1::mRuby::sun-1 3’UTR, cb-unc-119(+)] IVltIs37 [(pAA64) pie-1::mCherry::his-58 + unc-119(+)] IV*. SMW24 was also crossed with SMW22 to generate SMW33: *Pzyg-9::degron::EmGFP::zyg-9 (C6->T— PAM site mutation)* II; *unc-119(ed3)*; *ieSi38 [Psun-1::TIR1::mRuby::sun-1 3’UTR, cb-unc-119(+)] IV; weIs21 [pJA138 (Ppie-1::mCherry::tub::pie-1 3’UTR)]; unc-119(ed3) III*.

**Table 1.**
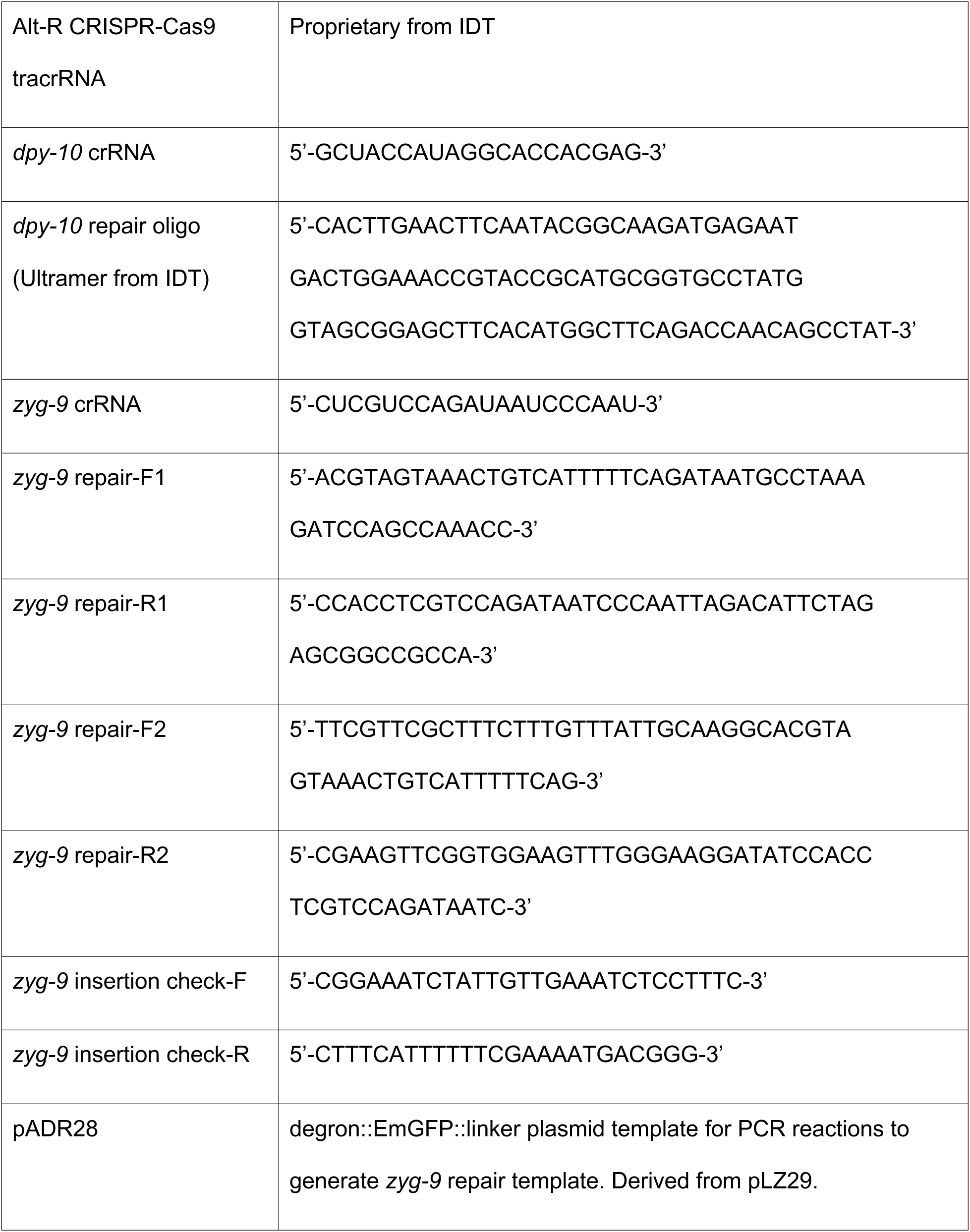
CRISPR/Cas9 Information.

### RNAi

RNAi was performed as in (Davis-Roca et al., 2018). Briefly, from a feeding library (Fraser et al., 2000; Kamath et al., 2003), individual RNAi clones were picked and grown overnight at 37°C in LB with 100μg/ml ampicillin. Overnight cultures were spun down and plated on NGM (nematode growth media) plates containing 100μg/ml ampicillin and 1mM IPTG. Plates were dried overnight. Worm strains were synchronized by bleaching gravid adults and letting the eggs hatch overnight without food. L1s were then plated on RNAi plates and grown to adulthood at 15°C for 5–6 days.

### Immunofluorescence and antibodies

Immunofluorescence was performed by dissecting worms into M9 or Meiosis Media (0.5 mg/mL Inulin from dahlia tubers (CAS Number 9005-80-5), 25 mM HEPES, pH 7.5, 60% Leibovitz’s L-15 Media (Gibco 11415-049), 20% Heat-inactivated Fetal Bovine Serum (Laband et al., 2018)), freeze cracking embryos, and plunging into -20°C methanol. Embryos were fixed for 35–45 minutes, rehydrated in PBS, and blocked in AbDil (PBS plus 4% BSA, 0.1% Triton X-100, 0.02% Na-Azide) for 30 minutes. Primary antibodies were incubated overnight at 4°C. The next day, embryos were washed 3x with PBST (PBS plus 0.1% Triton X-100), incubated in secondary antibody for 1 hour and 15 minutes, washed again as before, incubated in mouse anti-α-tubulin-FITC for 1.5 hours, washed again, and incubated in Hoechst (1:1000 in PBST) for 15 minutes. Embryos were then washed 2x with PBST, mounted in 0.5% p-phenylenediamine, 20mM Tris-Cl, pH 8.8, 90% glycerol, and sealed with nail polish; except for the overnight primary, the entire procedure was performed at room temperature. For experiments with staining of GFP::ZYG-9 (SMW24) with mouse anti-GFP, embryos were blocked in AbDil overnight at 4°C and incubated in primary antibody for 2 hours at room temperature. Primary antibodies used in this study: rabbit anti-ASPM-1 (1:5000, gift from Arshad Desai) (Wignall and Villeneuve, 2009), rat anti-KLP-18 (1:500, gift from Olaf Bossinger) (Segbert et al., 2003), rabbit anti-MESP-1 (1:3000) (Wolff et al., 2016), mouse anti-GFP (1:200; Invitrogen). Directly conjugated mouse anti-α-tubulin-FITC (DM1α, Sigma) and Alexa-fluor directly conjugated secondary antibodies (Invitrogen) were used at 1:500. All antibodies were diluted in AbDil.

### Generation of TAC-1 antibody

Based on previous predictions of TAC-1 domains and structure (Bellanger and Gonczy, 2003), we selected a 15aa sequence (PFNGSQNGHPENEEP) from the N-terminus of TAC-1; this region was chosen for relatively accessible residues as an epitope and due to this region’s lack of predicted interaction with other proteins. A rabbit polyclonal antibody was produced against this peptide sequence by ProteinTech, and the final antibody was affinity purified using a Sulfolink Immobilization Kit to generate the column. This antibody was validated for proper localization via IF imaging in both *control(RNAi)* and *tac-1(RNAi)* conditions. Due to relatively high background staining, this antibody was preabsorbed prior to use in immunofluorescence imaging. To preabsorb, ∼100 worms (from the same background strain as the experimental slides) were grown under *tac-1(RNAi)* conditions. These worms were dissected at adulthood and subjected to our standard fixation methods. After blocking, these worms were incubated with TAC-1 antibody (1:50 diluted in AbDil) and left overnight at 4°C. The following day, the TAC-1 solution was removed and stored for 24-48 hours at 4°C prior to being used in another set of IF slides containing the experimental conditions.

### Microscopy

All fixed imaging was performed on a DeltaVision Core deconvolution microscope with a 100x objective (NA = 1.4) (Applied Precision). This microscope is housed in the Northwestern University Biological Imaging Facility supported by the NU Office for Research. Image stacks were obtained at 0.2μm z-steps and deconvolved using SoftWoRx (Applied Precision). All images in this study were deconvolved and displayed as full maximum intensity projections of data stacks encompassing the entire spindle structure, unless stated otherwise.

### Time-lapse Imaging

Two-color live imaging was performed using a spinning disk confocal microscope with a 63x HC PL APO 1.40 NA objective lens. A spinning disk confocal unit (CSU-X1; Yokogawa Electric Corporation) attached to an inverted microscope (Leica DMI6000 SD) and a Spectral Applied Imaging laser merge ILE3030 and a back-thinned electron-multiplying charge-coupled device (EMCCD) camera (Photometrics Evolve 521 Delta) were used for image acquisition. The microscope and attached devices were controlled using Metamorph Image Series Environment software (Molecular Devices). For the acquisitions, 7 z-stacks at 2μm increments were taken every 30 seconds at room temperature. Images were processed using ImageJ. Images are shown as maximum intensity projections of the entire spindle structure. Live, intact worms were mounted on a homemade slide holder on 4% agarose in M9 pads that were made on Bob Dylan’s “A Shot of Love” vinyl LP (CBS) to make grooves for immobilizing worms (Rivera Gomez and Schvarzstein, 2018). The worms were picked into a drop of M9, 20μM serotonin creatinine sulfate monohydrate (Sigma), 2% tricaine, and 0.4% tetramisole and covered with a coverslip. The spinning disk microscope is housed in the Northwestern University Biological Imaging Facility supported by the NU Office for Research.

### Auxin Treatment

For long-term auxin treatment, worms were transferred onto NGM plates containing 1mM auxin (indol-3-acetic acid, Alfa Aesar) seeded with OP-50, incubated overnight, and then processed for immunofluorescence as described above.

For short-term auxin treatment for fixed imaging, whole worms were picked into a drop of L-15 blastomere media (0.5 mg/mL Inulin from dahlia tubers (CAS Number 9005-80-5), 25 mM HEPES, pH 7.5, 60% Leibovitz’s L-15 Media (Gibco 11415-049), 20% Heat-inactivated Fetal Bovine Serum) with 1mM auxin (indol-3-acetic acid, Alfa Aesar) and incubated for 25-30 minutes in a humidity chamber before dissection and freeze cracking. For vehicle treatment, 0.25% ethanol in L-15 blastomere media was used. The rest of the protocol is the same as the immunofluorescence procedure described above.

For acute auxin treatment of oocytes for live imaging, worms were picked into a drop of Meiosis media (described in the immunofluorescence section) containing either 100 µM auxin or vehicle (ethanol). The worms were quickly dissected to allow oocytes to enter the media, covered with a coverslip, sealed with Vaseline, and immediately imaged at room temperature using the spinning disk confocal described above. For more details on auxin protocols, see (Divekar et al., 2021a; Divekar et al., 2021b).

### Western Blotting

300 whole adult worms (per sample) were grown on control (empty vector) RNAi plates and treated in a manner identical to the auxin protocols utilized in this paper (30 minutes in auxin solution for short-term AID, 240 minutes on auxin plates for long-term AID). Worms were subsequently bleached to remove worm bodies and harvest embryos, and these samples were spun down (800 rcf for 1 minute), rinsed in M9 and spun down again, and then M9 was removed to leave only clean embryos. The pellet was then mixed with SDS lysis buffer, boiled for 10 minutes (using a heat block at 95°C), and run on a 4-20% gradient Tris-Glycine gel (BioRad Mini-PROTEAN TGX) at 80V for ∼1.5 hours. Protein was transferred onto nitrocellulose via a BioRad Trans-blot Turbo apparatus (semi-dry in a 10% MeOH, 25mM Tris, 192mM glycine transfer buffer) at 25V for 30 minutes. Blot was placed on rocker and blocked in 5% milk in TBS/0.1%Tween overnight at 4°C, separated into two pieces, and then incubated with indicated primary antibodies overnight at 4°C (1:1000 mouse anti-degron or 1:5000 mouse anti-tubulin). The entire blot was then washed with TBS/0.1% Tween, treated with anti-mouse HRP (1:5000) for 1.5 hours at RT, washed again, incubated for 4 minutes in BioRad Clarity ECL solution, and then exposed for 5 minutes.

### FRAP

FRAP experiments were performed on the same spinning disk confocal microscope mentioned above using a 63x HC PL APO 1.40 NA oil immersion objective. Photobleaching was performed with a 405 nm laser (5.5 mW-11.0 mW output). Poles were bleached using 5 repetitions of 100ms and images were taken at 5-10s intervals. Analysis of the recovery curves and the half-time recovery were carried out with FIJI to obtain raw fluorescence data and the curves were fit using a custom Python script (https://github.com/justinfinkle/mullen-frap). For all FRAP acquisitions except ASPM-1 meiotic spindles, oocytes and embryos were mounted as previously described (Laband et al., 2018). Briefly, oocytes and embryos were dissected into a drop of Meiosis media (0.5 mg/mL Inulin from dahlia tubers (CAS Number 9005-80-5), 25 mM HEPES, pH 7.5, 60% Leibovitz’s L-15 Media (Gibco 11415-049), 20% Heat-inactivated Fetal Bovine Serum) and mounted in a homemade slide mount. Bleaching of the mitotic centrosomes was done in the EMS cell in the 4-cell embryo, and MI and MII meiotic spindles were bleached for the acentrosomal spindle pole FRAP experiments. For ASPM-1 meiotic spindle FRAP experiments live, intact worms were mounted on 3-5% agarose, M9 pads in 50% live imaging solution (modified S-basal [50mM KH2PO4,10mM K-citrate, 0.1M NaCl, 0.025mg/ml cholesterol, 3mM MgSO4, 3mM CaCl2, 40mM serotonin creatinine sulfate monohydrate]), 50% 0.1 micron polystyrene Microspheres (Polysciences Inc.), and covered with a coverslip. Only data from bleached spindles that progressed to anaphase were included in our analysis.

## Data Analysis and Quantification

**Figure 1B:** Oocytes were imaged using the same set-up as described for the FRAP analysis. A 10 pixel wide x 12μm line profile analysis was performed in ImageJ on max projected images of 10 different metaphase spindles after background subtraction.

**Figure 1C:** IF images of oocyte spindles were used for linescan measurements of ASPM-1 and ZYG-9 channels. A 3 x 12µm line profile analysis was performed in ImageJ on max projected images of 23 different metaphase spindles after background subtraction.

**Figure 1D:** IF images of oocyte spindles stained for DNA, tubulin, ASPM-1, and ZYG-9 were scored as bipolar if ASPM-1 was enriched at the two poles, or multipolar/collapsed if ASPM-1 localized to multiple foci or was diffuse throughout the spindle. In oocytes from worms not treated with auxin, 1/34 spindles from 2 biological replicates were multipolar. In oocytes from worms plated on auxin-containing plates for 24 hours, 44/45 spindles from 2 biological replicates were multipolar or collapsed.

**Figure 1E:** IF images of 1-cell stage embryos were stained for DNA, tubulin, ASPM-1, and ZYG-9 were scored as aligned if the spindle was oriented parallel to the long axis of the embryo, or misaligned if the spindle was oriented perpendicular to this axis. In zygotes from worms not treated with auxin, 4/4 spindles from 2 biological replicates were aligned. In zygotes from worms plated on auxin-containing plates for 24 hours, 8/8 spindles from 2 biological replicates were misaligned.

**Figure 1 – figure supplement 1A:** 1A shows a representative movie of the *zyg-9(RNAi)* phenotype; we observed this phenotype in 3/3 movies, and a similar phenotype was reported in (Chuang et al., 2020).

**Figure 1 – figure supplement 1B:** ASPM-1 is enriched at spindle poles in control oocyte spindles (van der Voet et al., 2009). Following ZYG-9 depletion, ASPM-1 localized to multiple foci in 14/14 spindles, as shown in the representative image.

**Figure 1 – figure supplement 1C:** KLP-18 and MESP-1 are enriched at spindle poles in control oocyte spindles (Segbert et al., 2003; Wolff et al., 2016). Following ZYG-9 depletion, KLP-18 and MESP-1 co-localized at multiple foci in 18/18 spindles, as shown in the representative image.

**Figure 1 – figure supplement 1D:** Live, intact worms expressing GFP::tubulin, GFP::histone (EU1067) fed either control(RNAi) or zyg-9(RNAi)-expressing bacteria were anesthetized in 0.2% tricaine, 0.02% levamisole in M9 and viewed on a Leica DM5500B widefield fluorescence microscope. Spindles in embryos in the -1, spermatheca, and +1 positions within the gonad were scored for microtubule organization by eye. A spindle was scored as: ”cage” if microtubule bundles could be clearly seen after the haze of histone::GFP in the nucleus had disappeared, which indicates that the nuclear envelope had broken down; “multipolar” if it had prominent microtubule bundles that formed more than two organized poles; “collapsed” if the microtubule structure had collapsed around the chromosomes and lacked prominent bundles and organized poles; ”bipolar” if there were two organized spindle poles; ”anaphase” if there were two or more sets of segregated chromosomes. Control -1 (n=30): 36% cage (11/30), 13% NEBD No MTs (4/30), 43% multipolar (13/30), 3% bipolar (1/30), 3% collapsed (1/30). Control spermatheca (n=30): 87% multipolar (26/30), 13% bipolar (4/30). Control +1 (n=94): 10% multipolar (9/94), 43% bipolar (40/94), 9% collapsed (8/94), 38% anaphase (36/94). zyg-9(RNAi) -1 (n=34) 32% cage (11/34), 44% NEBD No MTs (15/34), 24% multipolar (8/34). zyg-9(RNAi) spermatheca (n=25): 92% multipolar (23/25), 4% bipolar (1/25), 4% collapsed (1/25); zyg-9(RNAi) +1 (n=191): 50% multipolar (96/191), 2% bipolar (4/191), 25% collapsed (48/191), 22.5% anaphase (43/191).

**Figure 1 – figure supplement 2:** For panel B, we captured multiple images of each stage, which all showed the localization pattern displayed in the representative images. The number of images captured were: 6 Pre-NEBD, 11 Cage/Multipolar, 9 Multipolar, 41 Bipolar, 16 Early anaphase, and 14 Late anaphase.

**Figure 2C:** Pole phenotypes were quantified by eye in unarrested *(control(RNAi))* and Metaphase I-arrested *(emb-30(RNAi))* oocytes in the presence of either vehicle or 1 mM auxin. Spindle poles were scored as fragmented if there were clear defects in spindle pole organization including excess tubulin and ASPM-1 signal in the area near the spindle poles and splaying or fragmentation of the tubulin and ASPM-1 signal at the poles. In unarrested conditions without auxin, 99/106 (93.4%) poles were focused while 7/106 (6.6%) were fragmented. With addition of auxin, we observed 13/79 (16.5%) poles were focused while 66/79 (83.5%) were fragmented. For metaphase-arrested conditions without auxin, 81/86 (94.2%) poles were focused while 5/86 (5.8%) were fragmented. With addition of auxin, we observed 27/90 (30%) poles were focused while 63/90 (70%) were fragmented.

**Figure 2D:** Midspindle phenotypes were quantified by eye in unarrested *(control(RNAi))* and Metaphase-I arrested *(emb-30(RNAi))* oocytes in the presence of either vehicle or 1 mM auxin. Midspindle microtubule splaying was determined by observing microtubule bundles near chromosomes. If microtubules could be seen splaying outwards into the cytoplasm, with no clear connection to another microtubule bundle or to the chromosomes, that spindle was considered splayed. . In unarrested conditions without auxin, 100/106 (94.3%) midzones were bundled while 6/106 (5.7%) were splayed. With addition of auxin, we observed 21/79 (26.6%) midzones were bundled while 58/79 (73.4%) were splayed. For metaphase-arrested conditions without auxin, 85/86 (98.8%) midzones were bundled while 1/86 (1.2%) were splayed. With addition of auxin, we observed 43/90 (47.8%) midzones were bundled while 47/90 (52.2%) were splayed.

**Figure 2E:** Imaris 3D Imaging Software (Bitplane) was used for spindle length measurements (procedure modified from (Davis-Roca et al., 2017; Heath and Wignall, 2019)). To calculate spindle length, the “Surfaces” tool was first used to determine the volume of each pole stained with ASPM-1 and then to assign the center of the volume for each pole. The distance between these two center points was then measured as the spindle length.

**Figure 3C:** Oocyte spindles were quantified using the FIJI plugin Coloc2 to determine the Pearson coefficient between the ZYG-9 and ASPM-1 channels. All r values were placed into boxplots using Rstudio (boxes represent first quartile, median, and third quartile); the mean values of the Pearson coefficient in *control(RNAi)* and *tac-1(RNAi)* conditions were compared against each other using a two-tailed t-test. Images for each condition were collected from at least three biological replicates, and the number of images for each condition was listed above the boxplots.

**Figure 3D:** Oocyte spindles were quantified by eye as described for Figure 2C. Exact percentages for *control(RNAi)* were 95.1% (78/82) focused, 4.9% (4/82) fragmented (No Auxin), and 18.1% (17/94) focused, 81.9% (77/94) fragmented (With Auxin). Exact percentages for *tac-1(RNAi)* were 13.9% (5/36) focused, 86.1% (31/36) fragmented (No Auxin) and 11.1% (3/27) focused, 88.9% (24/27) fragmented (With Auxin). Images for each condition were collected from at least three biological replicates.

**Figure 3E:** Oocyte spindles were quantified by eye as described for Figure 2D. Exact percentages for *control(RNAi)* were 91.4% (75/82) bundled, 8.6% (7/82) splayed (No Auxin) and 22.3% (21/94) bundled, 77.7% (73/94) splayed (With Auxin). Exact percentages for *tac-1(RNAi)* were 38.9% (14/36) bundled, 61.1% (22/36) splayed (No Auxin) and 29.6% (8/27) bundled, 70.4% (19/27) splayed (With Auxin). Images for each condition were collected from at least three biological replicates.

**Figure 4 – figure supplement 1:** Verification of antibody staining was done by comparing *control(RNAi)* and *tac-1(RNAi)* worms. Localization of TAC-1 and ZYG-9 to centrosomes was consistent across 5/5 *control(RNAi)* embryos and loss of TAC-1 and ZYG-9 at centrosomes was observed in 5/5 *tac-1(RNAi)* embryos.

**Figure 5B:** The volume of the spindle pole was determined using ASPM-1 staining as described in (Hollis et al., 2020). Briefly, Imaris 3D Imaging Software (Bitplane) was used for spindle pole volume measurements. The “Surfaces” tool was used to determine the volume of each pole, using ASPM-1 staining to define the pole. Images for each condition were collected from at least three biological replicates.

**Figure 5C:** 5C shows representative movies of embryos dissected into either EtOH (vehicle), or auxin (to deplete ZYG-9). Following auxin treatment, monopolar spindle organization was not disrupted upon ZYG-9 depletion in 5/5 movies.

## Statistical methods

All statistical analysis was done using a student’s two-tailed t test. Data distribution was assumed to be normal, but this was not formally tested.

## ACKNOWLEDGMENTS

We thank all members of the Wignall lab for support and discussion, and Hannah Horton and Claire Strothman for critical reading of the manuscript. We thank Bruce Bowerman, Olaf Bossinger, Arshad Desai, and Abby Dernburg for antibodies and strains. Some strains were provided by the *Caenorhabditis* Genetics Center (CGC), which is funded by NIH Office of Research Infrastructure Programs (P40 OD010440). This work was funded by NIH R01GM124354 (to SMW), by the Carcinogenesis Training Grant T32 CA009560 (to TJM and GCM), by the Cell and Molecular Basis of Disease Training grant T32GM008061 (to ERC), by American Heart Association Predoctoral Fellowship 17PRE33440016 (to IDW), and NIH/NIGMS Molecular Biophysics Training Grant T32GM008382 (to IDW).

---

**Figure 1 – figure supplement 1:**
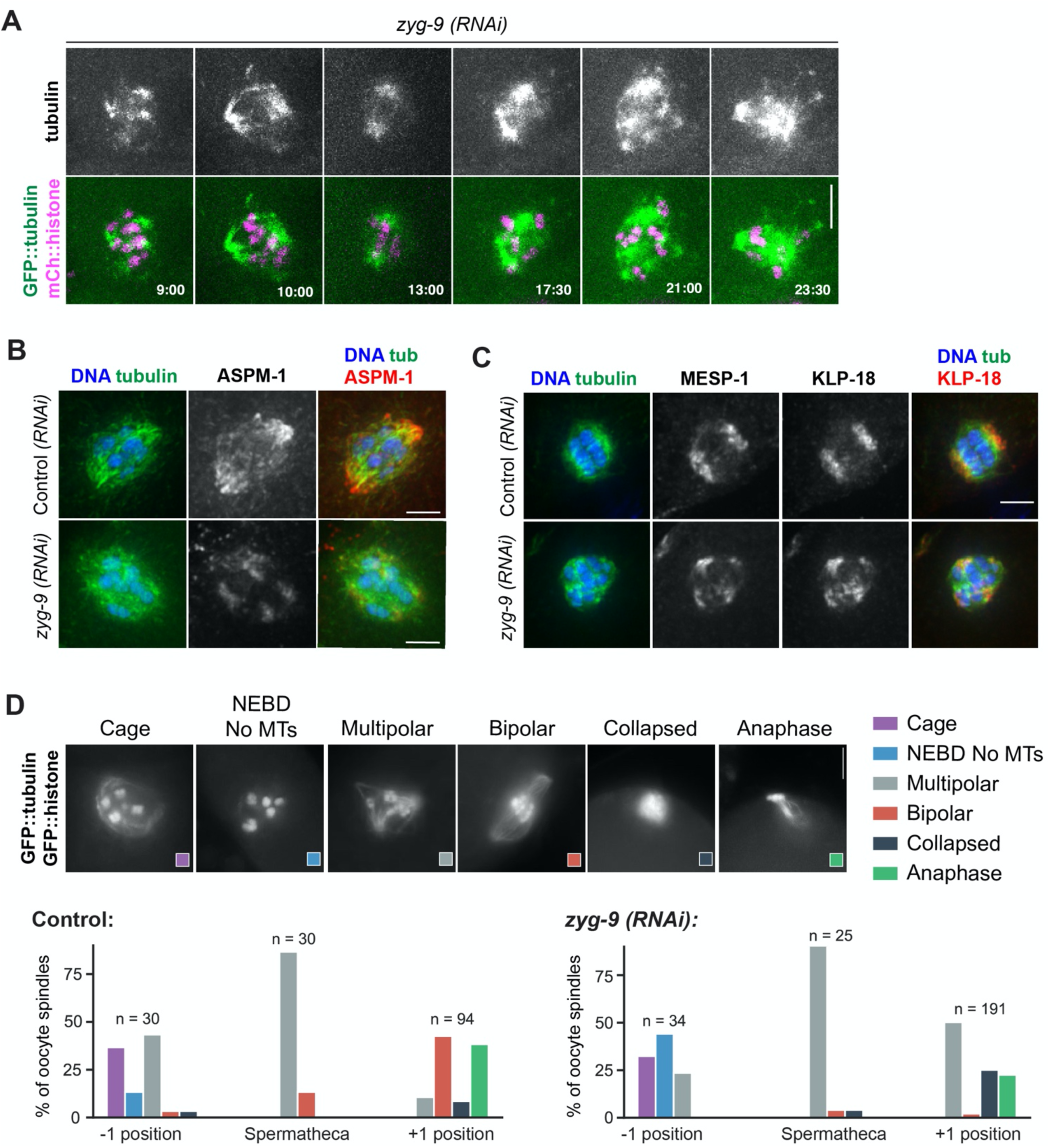
ZYG-9 is required for acentrosomal spindle assembly and is enriched at spindle poles. (A) Movie stills from *zyg-9(RNAi)* oocytes expressing GFP::tubulin (green) and mCherry::histone (magenta). Following *zyg-9(RNAi)*, microtubules appear to nucleate in close proximity to the chromosomes (9:00). As spindle assembly progresses, a bipolar structure transiently forms (13:00) before the poles begin to fragment into multiple pole-like foci (21:00). This multipolar structure then collapses into a mass of tubulin surrounding the chromosomes (23:30). Bar = 5µm. Timestamp = mm:ss. (B) Control and *zyg-9(RNAi)* spindles stained for DNA (blue), tubulin (green), and ASPM-1 (red). In control spindles, ASPM-1 marks the two poles of the bipolar spindle. Following *zyg-9(RNAi)*, ASPM-1 stains multiple, discrete structures within the oocyte spindle. Bar = 2.5µm. (C) Control and *zyg-9(RNAi)* spindles stained for DNA (blue), tubulin (green), MESP-1 (not in merge), and KLP-18 (red). In control spindles, MESP-1 and KLP-18 colocalize at the two poles of the bipolar spindle. Following *zyg-9(RNAi)*, MESP-1 and KLP-18 stain multiple, discrete structures within the oocyte spindle. Bar = 2.5µm. (D) The organization of the *C. elegans* germ line allows extrapolation of temporal information based on where the oocyte is located. Oocytes in the -1 position are ovulated into the spermatheca where they are fertilized and begin the meiotic divisions. The fertilized oocytes exit the spermatheca to the +1 position where they continue meiosis before beginning the mitotic divisions. Quantification of spindle phenotypes in control (left), *zyg-9(RNAi)* (right) , and examples of phenotypes (top). This analysis was performed by viewing spindles in live worms expressing GFP::tubulin; GFP::histone. In *zyg-9(RNAi)* oocytes, there is an increase in the number of multipolar and collapsed spindles and a decrease in the number of bipolar spindles found in the +1 position. n represents the number of spindles analyzed at each position. Bar = 5µm.

**Figure 1 – figure supplement 2:**
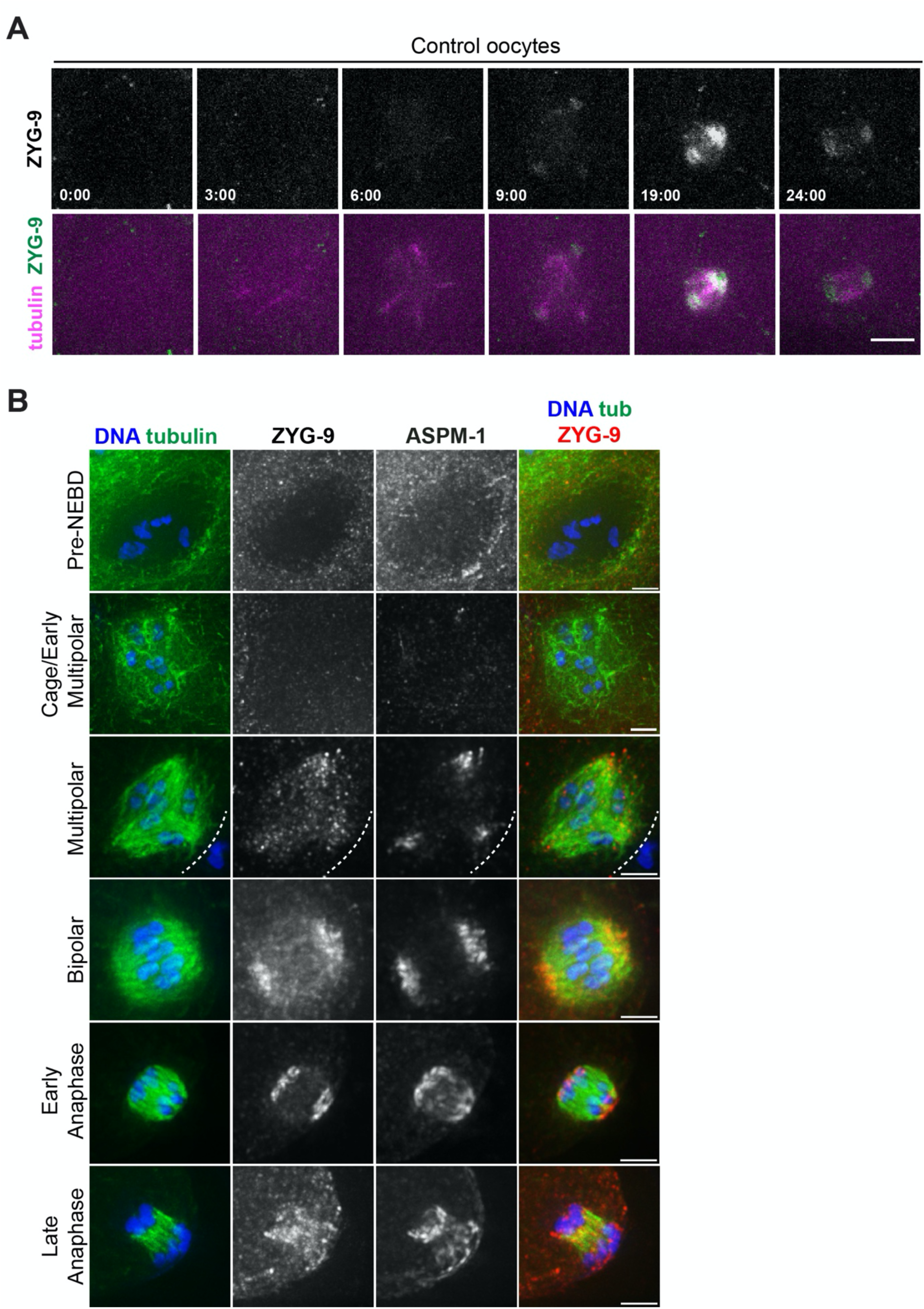
Imaging of degron::GFP::ZYG-9 during spindle assembly. (A) Movie stills from an oocyte expressing degron::GFP::ZYG-9 (green) and mCherry::tubulin (magenta). ZYG-9 initially localizes to the spindle microtubules at the multipolar stage (6:00) and becomes progressively enriched on the spindle poles as meiosis progresses (9:00 - 19:00) before dissociating during anaphase (24:00). Bar = 5µm. Timestamp = min:sec. (B) Oocyte spindles stained for DNA (blue), tubulin (green), ZYG-9 (red in merge), and ASPM-1. ZYG-9 localizes to spindle microtubules at the multipolar stage, becomes enriched at the spindle poles as spindle assembly proceeds, then begins to lose enrichment at the poles during anaphase. Bars = 2.5µm.

**Figure 1 – figure supplement 3:**
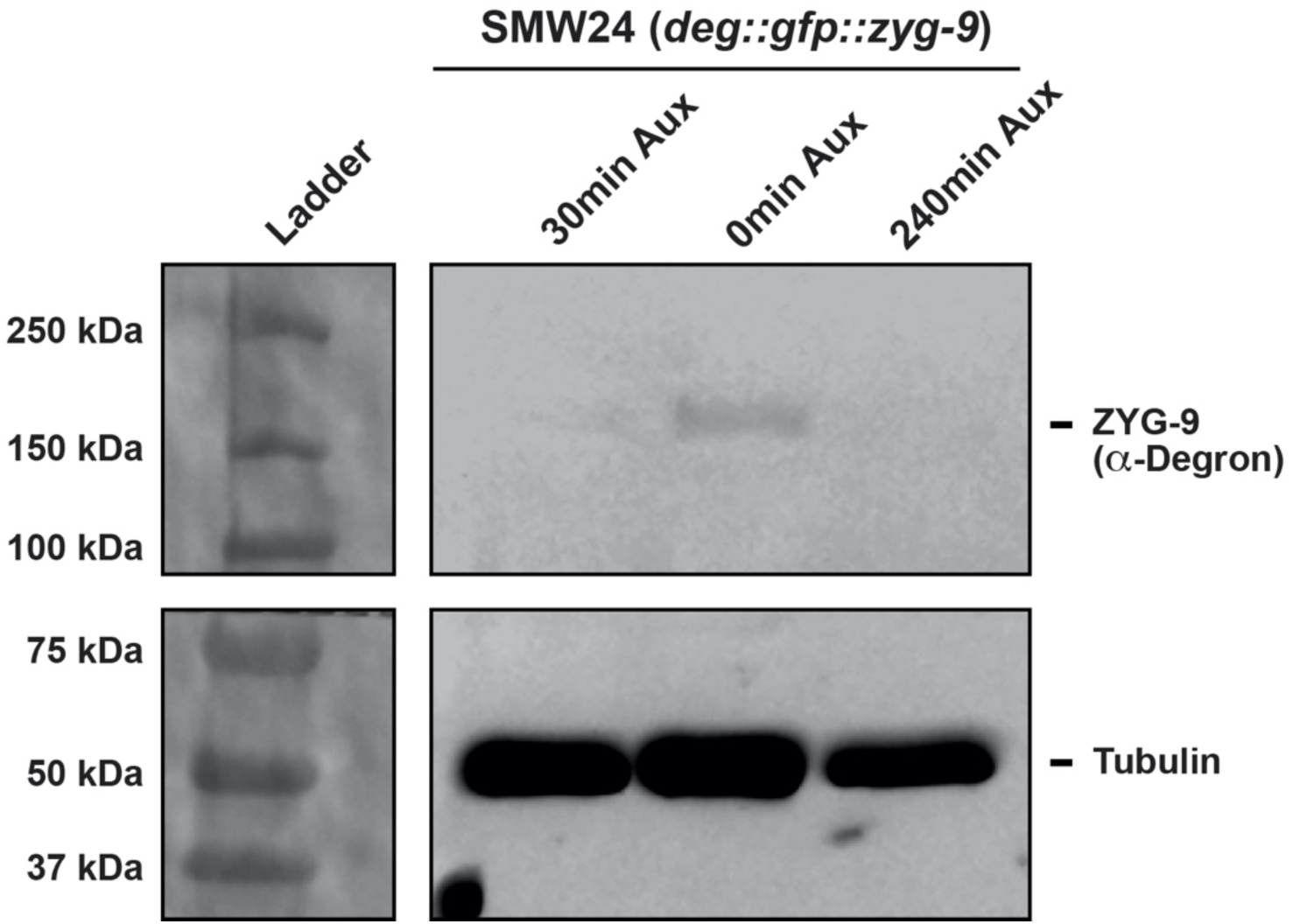
Analysis of ZYG-9 depletion using Western blotting. An embryo-only Western blot demonstrating the effectiveness of both short-term and long-term AID depletion of ZYG-9 in the ZYG-9-AID strain expressing degron::GFP::ZYG-9 (predicted to be ∼188 kDa). A faint band of ZYG-9 is present in control sample (middle lane), while no band is present in either the short-term AID (30 minute treatment via soaking; left lane) or long-term AID (240 minute treatment on plates; right lane) embryos, validating the efficiency of AID depletion. Tubulin was utilized as a loading control and ZYG-9 was detected with an anti-degron antibody.

**Figure 2 – figure supplement 1:**
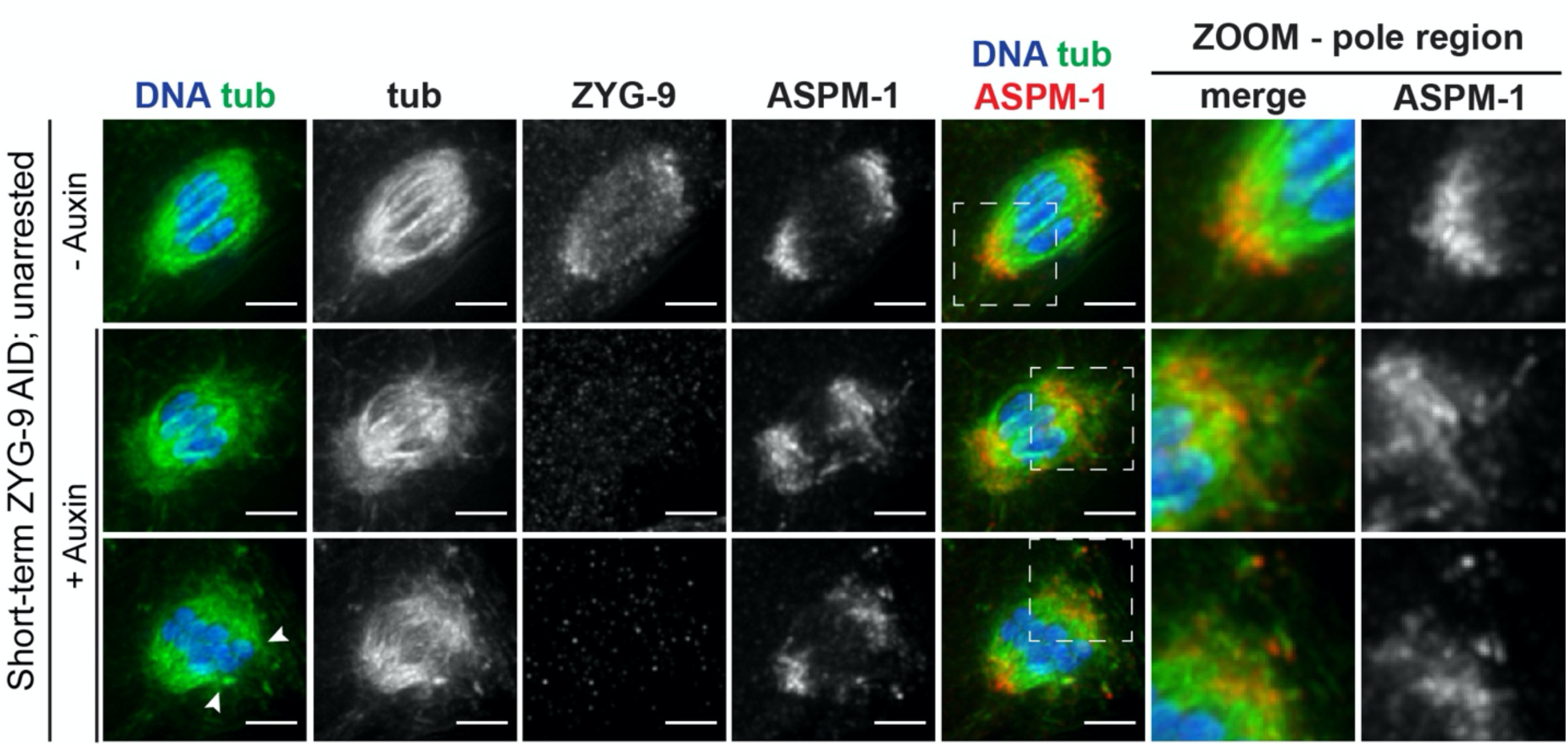
Short-term AID of ZYG-9 on unarrested bipolar spindles. IF imaging of oocyte spindles in *control(RNAi)* conditions treated with either vehicle or 1mM auxin solution for 25-30 minutes prior to dissection and fixation, stained for DNA (blue), tubulin (green), ZYG-9 (not in merge), and ASPM-1 (red). All ZYG-9 depletion phenotypes observed in Metaphase I-arrested conditions (Figure 2B) are apparent in unarrested conditions as well (midspindle disruption highlighted with arrowheads). Zooms of pole regions in auxin-treated spindles clearly display fragmentation of poles seen by dispersal of tubulin and ASPM-1 signal and individual fragments. Bars = 2.5 µm.

**Figure 4 – figure supplement 1:**
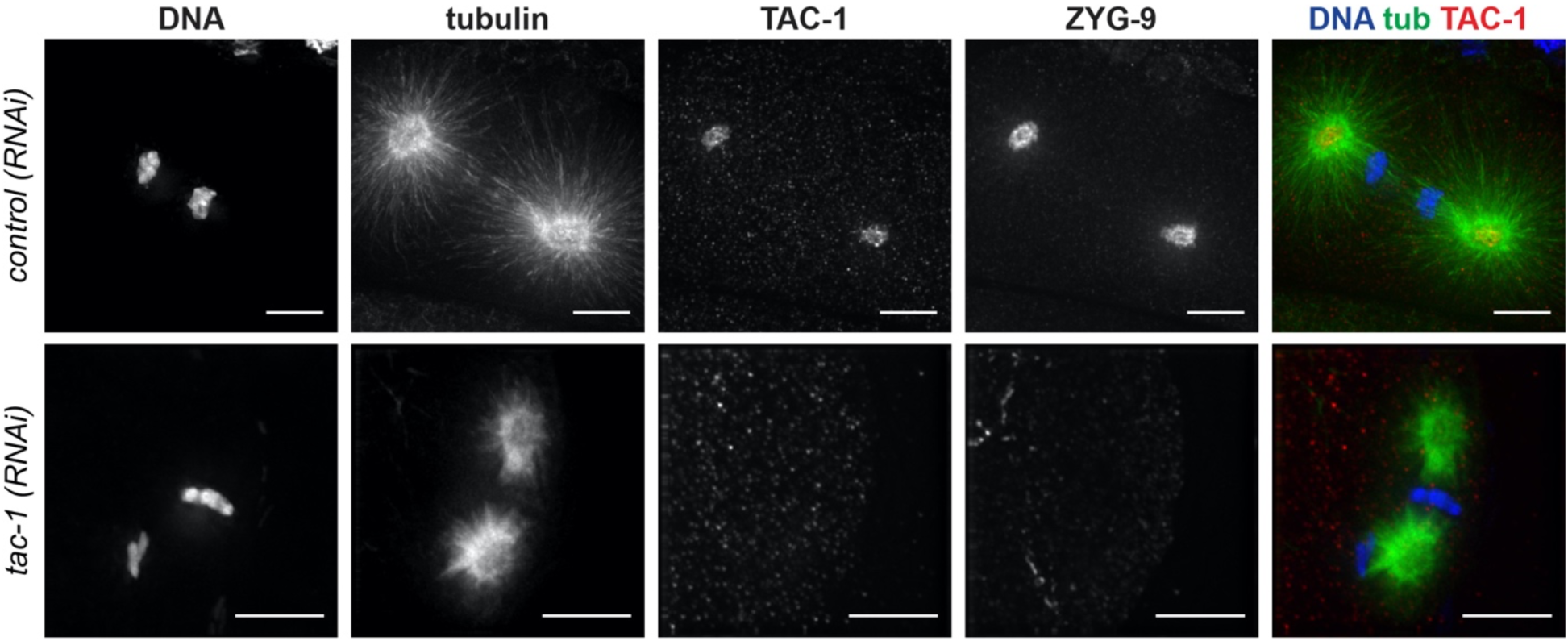
TAC-1 Antibody Validation. Verification of TAC-1 antibody using IF imaging of one-cell mitotic embryos. TAC-1 localizes to centrosomes, and colocalizes with ZYG-9, as previously described (Bellanger and Gonczy, 2003; Srayko et al., 2003). Utilizing *tac-1(RNAi)* demonstrates that TAC-1 staining is specific, as no staining occurs when TAC-1 is depleted. Also consistent with previous studies, loss of TAC-1 leads to loss of ZYG-9 at centrosomes and results in defects in mitotic spindle positioning and spindle length. Bars = 5µm.

**Figure 6 – figure supplement 1:**
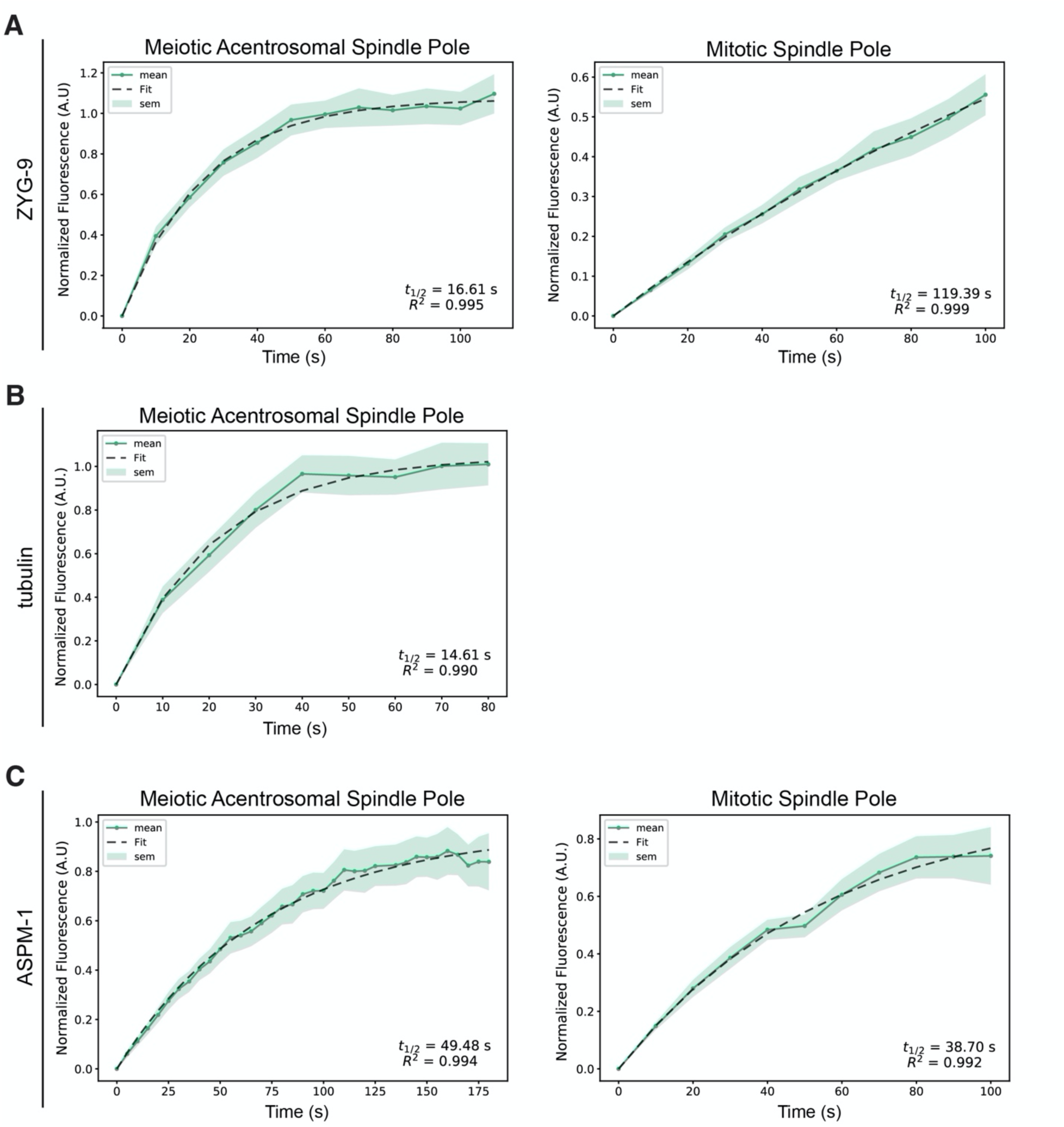
FRAP analysis with fit curves. (A-C) graphs of recovery curves and fit curves from the bleached poles of the FRAP experiments described in Figure 6. The mean is a solid line, the standard error of the mean is the shaded region, and the fit from the single exponential function is a dashed line. The t1/2’s were calculated from these fit curves.

## SUPPLEMENTAL VIDEOS

**Video 1. Live imaging of spindle assembly following *zyg-9(RNAi)*.**

*In utero* imaging of a *zyg-9(RNAi)* embryo expressing mCherry::histone and GFP::tubulin (left); GFP::tubulin alone is shown on the right. Corresponds to Figure 1 – figure supplement 1A. Following ZYG-9 depletion, a transient bipolar spindle forms but then poles split apart and become unstable. Time elapsed shown in (min):(sec). Scale bar = 10μm.

**Video 2. ZYG-9 localization during oocyte meiosis.**

*In utero* imaging of control embryo expressing mCherry::tubulin and GFP::ZYG-9 (left); GFP::ZYG-9 alone is shown on the right. Corresponds to Figure 1 – figure supplement 2A. ZYG-9 localizes to spindles at the multipolar stage, and is enriched at poles in metaphase and anaphase. Time elapsed shown in (min):(sec). Scale bar = 10μm.

**Video 3. Auxin treatment rapidly depletes ZYG-9 and causes spindle instability.**

Shows a metaphase-arrested embryo expressing mCherry::tubulin (left), dissected into Meiosis Media containing 100μM auxin to deplete ZYG-9. Corresponds to Figure 2A. Once dissected into auxin solution, rapid depletion of degron::GFP::ZYG-9 (right) is evident and spindle poles become unstable. Time elapsed shown in (min):(sec). Scale bar = 5μm.

**Video 4. Auxin treatment rapidly depletes ZYG-9 and causes spindle instability.**

Another example of a metaphase-arrested embryo expressing mCherry::tubulin (top), dissected into Meiosis Media containing 100μM auxin to deplete ZYG-9. Once dissected into auxin solution, rapid depletion of degron::GFP::ZYG-9 (bottom) is evident and spindle poles become unstable. Time elapsed shown in (min):(sec). Scale bar = 5μm.

**Video 5. Metaphase-arrested spindles remain stable without auxin treatment.**

Shows a metaphase-arrested embryo expressing mCherry::tubulin (left) and degron::GFP::ZYG-9 (right), dissected into a control Meiosis Media solution. Corresponds to Figure 2A. No major changes in spindle length or shape occur. Note that the spindle rotates end-on for a portion of the video, but when it rotates back it is clear that the morphology of spindle has not changed. Time elapsed shown in (min):(sec). Scale bar = 5μm.

**Video 6. Monopolar spindle organization does not change following acute ZYG-9 depletion.**

Shows an embryo expressing mCherry::tubulin (left) and degron::GFP::ZYG-9 (right) following *klp-18(RNAi)*, dissected into Meiosis Media containing 100μM auxin to deplete ZYG-9. Corresponds to Figure 5C. Upon dissection into auxin solution, ZYG-9 is rapidly depleted but the organization of the monopole does not noticeably change. Time elapsed shown in (min):(sec). Scale bar = 5μm.

**Video 7. Monopolar spindles remain stable without auxin treatment.**

Shows a *klp-18(RNAi)* embryo expressing mCherry::tubulin (left) and degron::GFP::ZYG-9 (right), dissected into a control Meiosis Media solution. Corresponds to Figure 5C. No major changes in monopole organization occurs during the course of the movie. Time elapsed shown in (min):(sec). Scale bar = 5μm.

**Video 8. FRAP of ZYG-9 at the poles of acentrosomal oocyte spindles.**

Shows a Metaphase II oocyte spindle labeled with degron::GFP::ZYG-9 and mCherry::histone; the polar body is on the left side of the spindle. During the video the pole on the left is photobleached but fluorescence quickly recovers. Corresponds to Figure 6A. Time elapsed shown in seconds. Scale bar = 2.5μm.

**Video 9. FRAP of ZYG-9 at the poles of mitotic centrosome-containing spindles.**

Shows a mitotic spindle in the EMS cell of the 4-cell embryo, labeled with degron::GFP::ZYG-9 and mCherry::histone. During the video the pole on the right is photobleached. Fluorescence recovers but more slowly than on acentrosomal poles. Corresponds to Figure 6A. Time elapsed shown in seconds. Scale bar = 5μm.

**Video 10. FRAP of tubulin at the poles of acentrosomal oocyte spindles.**

Shows a Metaphase II oocyte spindle labeled with degron::GFP::ZYG-9 and mCherry::histone; the polar body is on the right side of the spindle. During the video the pole on the bottom is photobleached but fluorescence quickly recovers. Corresponds to Figure 6B. Time elapsed shown in seconds. Scale bar = 2.5μm.

**Video 11′. FRAP of ASPM-1 at the poles of acentrosomal oocyte spindles.**

Shows a Metaphase II oocyte spindle labeled with GFP::ASPM-1 and mCherry::histone; the polar body is not visible in the selected z-stacks. During the video the pole on the left is photobleached. Corresponds to Figure 6C. Time elapsed shown in seconds. Scale bar = 2.5μm.

**Video 12. FRAP of ASPM-1 at the poles of mitotic centrosome-containing spindles.**

Shows a mitotic spindle in the EMS cell of the 4-cell embryo, labeled with GFP::ASPM-1 and mCherry::histone. During the video the pole on the left is photobleached. Corresponds to Figure 6C. Time elapsed shown in seconds. Scale bar = 5μm.

## SOURCE DATA LEGENDS

**Figure 1.** The source data for Figures 1B and 1C are provided. For Figure 1B, the x-axis position value is listed alongside the fluorescence intensity measurements for each file; the DNA measurements and ZYG-9 measurements are listed in separate tabs. The data for Figure 1C is listed in a third tab; for each file, the raw intensity values for ZYG-9 and ASPM-1 are provided for each position along the x-axis. Average values and standard errors are also listed.

**Figure 2.** The source data for Figure 2E is provided; measurements for unarrested *(control(RNAi))* and Metaphase I-arrested *(emb-30(RNAi))* spindles are listed in separate tabs. The spindle length measurements (in μm) are provided for each image, as well as the averages lengths and standard errors with and without auxin.

**Figure 3.** The source data for Figure 3C is provided; *control(RNAi)* and *tac-1(RNAi)* data are listed in separate tabs. The Pearson’s coefficient between ASPM-1 and ZYG-9 are provided for each of the measured images.

**Figure 5.** The source data for Figure 5B is provided. Volume measurements for control (minus auxin) and plus auxin are listed in separate tabs.

**Figure 6.** The source data for all graphs in Figure 6 are provided as separate tabs. The time (in seconds) is listed, along with the fluorescence intensity measurement at that timepoint for each of the analyzed movies; t=0 was set as the time of bleaching. The measurements for the bleached pole and the unbleached pole are listed in separate tabs.

## Notes

### Competing Interest Statement

The authors have declared no competing interest.

